# Functional, Biotinylproteomic and Bioinformatic Analysis of Both Cytoskeletal and Plastoskeletal Proteins in Plant Mechanoresponse

**DOI:** 10.1101/2024.11.30.626153

**Authors:** Kebin Wu, Nan Yang, Jia Ren, Shichang Liu, Kai Wang, Shuaijian Dai, Yinglin Lu, Yuxing An, Fuyun Tian, Zhaobing Gao, Zhu Yang, Yage Zhang, Weichuan Yu, Ning Li

**Author notes:** Co-first author (Kebin Wu, Nan Yang). Correspondence Address: Division of Life Science, The Hong Kong University of Science and Technology, Hong Kong, SAR, China.

## Abstract

To investigate the early signaling components of skeletal proteins in mediating Arabidopsis thigmomorphogenesis, both microscopic and proximity labeling (PL)-based quantitative biotinylproteomics were applied to investigate the subcellular location and putative interactors of a touch-responsive WPRa4 protein. These experiments have demonstrated that the cytoskeletal protein WPRa4 is localized nearby the plastid. Several cytosolic Plastid Movement-Impaired (PMI) proteins and a member of the plastidic translocon were identified as putative interactors of WPRa4, suggesting an integrated network of skeletal proteins linking the cytoskeleton with the plastid membrane. Further bioinformatic analysis of both Proximity Labeling- and XL-MS-based proteomic results suggested that Plastid Movement-Impaired 4 (PMI4) protein may serve as a candidate in mediating the plant touch response. The loss-of-function *pmi4* mutant showed neither the touch-induced bolting delay nor the rosette size reduction upon repetitive touches, suggesting that *pmi4* is a unique type of mutant of Arabidopsis thigmomorphogenesis. Moreover, the null mutant *pmi4* displayed a severe defect in the touch-induced Ca^2+^ oscillation. Further transcriptomic analysis performed on both the wild-type Arabidopsis and *pmi4* mutant indicated that the mutated *pmi4* gene suppressed the expression of a number of touch rapidly induced transcripts and a JA-responsive gene, *LOX2*. These findings led us to propose a revised touch force-sensing theory, in which the interconnected cytosolic and plastidic skeletal proteins serve as the early mechano-sensing components mediating Arabidopsis thigmomorphogenesis and the retrograde calcium signaling in response to touch.

## Introduction

It is widely acknowledged that cells can sense and respond to mechanical signals of their extracellular environment *via* numerous structures and signaling components, adapting to the surrounding conditions in a process known as mechanotransduction (1). Many components have been proven to facilitate this process, including integrins, stretch-active ion channels, receptor-like kinases, cytoskeletal filaments, *etc* (2). For example, mammalian cells utilize integrin to connect the extracellular matrix (ECM) with the F-actin cytoskeleton for mechanotransduction which enables cell spreading and migration (3). Furthermore, cellular intermediate filaments create a mechanical continuum that extends throughout the cell, linking the cell surface to the nucleus through transmembrane protein complexes in both the plasma and nuclear membranes (4). This specialized structure transmits extracellular forces into the cell, offering mechanical signals that affect signaling events and cell development (2).

While plants and animals might share certain common mechanisms underlying mechanotransduction despite their distant evolutionary origins, each species also possesses its own distinct mechanisms for detecting and responding to mechanical forces (5). Several hypotheses have been postulated in the past years for the plant force-sensing mechanisms, which include the theories of stretch-activated channels (6), intracellular turgor pressure and subsequent cytoplasmic streaming (7), as well as the plant-specific cell wall-plasma membrane-actin cytoskeleton continuum (7, 8), where the importance of the cytoskeleton in mechanotransduction is emphasized. Another major difference in mechanotransduction between plants and animals is the involvement of the plant-specific organelle, the plastid, whose role in gravity-sensing has been well-established based on the starch-statolith hypothesis (5). In Arabidopsis, it has been well-documented that the specialized type of sensory plastids that has a unique *MutS Homolog 1* (*MSH1*) genetic variation is able to mediate the systemic stress response (9). Furthermore, the epidermal cell-specialized plastids have been reported to participate in the plant immunity-related processes through plastid movement-related protein components, such as Chloroplast Unusual Positioning 1 (CHUP1) and J-domain Protein Required for Chloroplast Accumulation Response 1 (JAC1) (10). It is therefore proposed that both plastid-related and plastid movement-related proteins might function as the early force signaling components mediating the touch response in known pathways, such as calcium oscillation (11), post-translational modification of signaling component (12), gene expression (13), and phytohormone biosynthesis (14, 15).

To reveal the early protein signaling components of plant mechano-response, our laboratory has performed a functional phosphoproteomic study on Arabidopsis touch response, from which 24 phosphoprotein groups were found to be force-responsive within 40 seconds (12). These phosphoproteins include two groups of kinases, Mitogen-activated protein Kinase Kinase 1/2 (MKK1/MKK2) and Calcium-dependent Protein Kinases (CPKs), as well as a cytoskeletal structure component WPRa4 (12), which is a member of WEB1/PMI2-Related (WPR) protein family (16). Molecular genetic studies on both touch-responsive MKK1/2 kinases as well as the cytoskeletal protein WPRa4 revealed the regulatory functions of these proteins in Arabidopsis touch responses, especially in bolting delay (12, 17). In the present study, to address the role of a cytosol-localized plastid movement-related WPR family protein WPRa4 in plant mechano-sensory process, we again adopted a combinatorial Omics approach comprising of phenomics, biotinylproteomics, and bioinformatics. Both microscopic and *in planta* proximity-labeling (PL)-based (18) quantitative biotinylproteomic studies indeed revealed that WPRa4 is localized near the plastid.

A thorough bioinformatic investigation of proteomics results and literature indicated a possible role for a plastoskeletal (plastid skeleton) protein in plant touch response, this Plastid Movement-Impaired 4 (PMI4), or called Filamentous Temperature-Sensitive Z1 (FtsZ1) protein (19, 20), was therefore chosen as a candidate for the touch response study. At the same time, the Accumulation and Replication of Chloroplasts 3 (ARC3) protein, which also plays a role in the plastoskeleton Z-ring assembly and the mutant of which exhibits overly enlarged plastids and reduced plastid number (21), was selected as a control for Arabidopsis touch response studies. Surprisingly, only *PMI4* gene alone plays an important role in plant touch response. Especially, it severely suppressed both the touch-responsive retrograde Ca^2+^ oscillation and the touch-induced gene expression. Thus, these experimental results have demonstrated that both cytoskeletal and plastoskeletal proteins participate in the early force signaling during the thigmomorphogenesis of Arabidopsis. A novel mechano-sensing mechanism is hereby proposed to involve the plastidic gene PMI4-regulated retrograde calcium transient in the plant touch response.

### Experimental Procedures

#### Experimental Design and Statistical Rationale

The MS sample preparation follows the 4C workflow as described in Supplemental Fig. S2. In short, *in planta* TurboID-based proximity labeling (PL) of proximal protein with biotin on free lysine site (22) and *in vitro* dimethyl labelling of digested peptide (23) with C_12_H_2_ formaldehyde (light, +28.031 Da) or C_13_D_2_ formaldehyde (heavy, +34.063 Da) were employed as the first step (Supplemental Fig. S2, part 1). Light labeled peptide from TurboID/Col-0 was mixed with heavy labeled peptide from WPRa4-TID/wpra4 as the forward sample, and heavy labeled peptide from TurboID/Col-0 was mixed with light labeled peptide from WPRa4-TID/wpra4 as the reciprocal sample. In total, three biological replicates were conducted for each of the Forward (F1, F2, F3) and Reciprocal (R1, R2, R3) replicates, yielding six high purity biotinylated peptide samples for MS analysis (Supplemental Fig. S2, part 2). The six MS data sets obtained from Mascot searches for both heavy and light dimethyl-labeled biotinylpeptide pairs were summarized in Supplemental Table S1. Extracted ion chromatography (XIC, MS1 ion intensity)-based quantitative biotinylproteomics was achieved by in-house built software SQUA(23) (Supplemental Fig. S2, part 3). The quantifiable biotinylated proteins were described in Supplemental Table S2 with 21 putative interactors of WPRa4 of both XIC ion intensity Log_2_ ratio > 2.0 and Log_2_(BOR) > 0. Using GO enrichment on those 21 putative interactors, the role of plastid in thigmomorphogenesis was studied along with other functional validation (Supplemental Fig. S2, part 4).

In the SQUA-based quantitation, the identification of both light and heavy labeled biotinylpeptides was accomplished using the Mascot search engine, maintaining a false discovery rate (FDR) of 1%(22). The log2-ratios of biotinylpeptide MS1 ion intensities were initially adjusted based on either a three-way analysis of variance (ANOVA) or the empirical Bayes method, depending on the observed batch effects (23). The XIC quantitation selection of putative interactors of WPRa4 was conducted using a one-sample t-test on the log2-ratios of biotinylpeptide ion intensities on MS1 level. The p-values from the t-test on the log2-ratios of Unique Peptide Arrays (UPAs) of biotinylpeptides were subjected to multiple hypothesis testing correction via the Benjamini and Hochberg (BH) procedure (24), with a threshold set at a 5% false discovery rate (BH-FDR). Biotin Occupancy Ratio (BOR) (22) was also adapted in the putative interactors selection in this study.

#### Plant materials and genotypes

Arabidopsis wild type *Columbia-0* (*Col-0*) and T-DNA insertional mutant line SALK_023690 (T-DNA insertional mutagenized AT5G55860, named as *treph1*-*1* as in ref. (12)), as *wpra4* herein were obtained from the *Arabidopsis* Biological Resource Center (ABRC, Columbus, OH, USA). T-DNA insertional mutant lines SALK_073878 (T-DNA insertional mutagenized AT5G55280, *ftsz1/pmi4* referred to as *pmi4* in this study), SALK_057144 (T-DNA insertional mutagenized AT1G75010, referred as *arc3* in this study) and CS68684 (Double mutant generated by crossing *pmi4* (SALK_073878) and *arc3* (SALK_057144)) were a gift from Professor Osteryoung KW. The T-DNA left border (LB1) primer, *pmi4, and ARC3* gene-specific left and right primers (LP, RP), were synthesized to confirm the insertional mutagenesis as described before in (ref. (12)). (Genotyping information was shown in Supplemental Fig. S7A-B). The *wpra4* line was used as the transgenic background of *WPRa4-TID*. *Col-0, pmi4, arc3, and arc3pmi4* mutants were used for the automated hair touch experiment. *Col-0* and *pmi4* mutants were used for transcriptomic analysis experiments under the automated hair touch treatment.

For the cellular Ca^2+^ spike monitoring experiment, we introduced the CRISPR-Cas9 editing system (Genovo Bio, Tianjin, China) (25) into the *Pro35S::Aequorin/Col-0* transgenic plant (named as *AEQ/Col-0*, a gift from Dr. Knight) (11), yielded *AEQ*/*pmi4*-*c1* (*AEQ*/*pmi4-c*), *AEQ*/*pmi4-c2*, *AEQ*/*arc3-c1* (*AEQ*/*arc3-c*), *AEQ*/*arc3pmi4-c1* (*AEQ*/*arc3 pmi4-c*), *and AEQ*/ *piezo1-c1* (*AEQ*/*piezo1-c*) mutants (Supplemental Fig. S12A). T3 generation plants were sequenced as homozygous mutants and then selected for experiment. These CRISPR-Cas9-generated mutants were used in the Ca^2+^ spike monitoring experiments. *AEQ/Col-0*, *AEQ*/*pmi4-c*, *AEQ*/*arc3-c*, and *AEQ*/*arc3pmi4-c* plants were applied to test the touch-induced Ca^2+^ oscillation as shown in Fig. 5. The results of touch-induced Ca^2+^ oscillation of *AEQ*/*pmi4-c2*, and *AEQ*/*piezo1-c* plants were shown in Supplemental Fig. S12B.

All plant growth medium components and chemicals were purchased from Sigma if not specified otherwise. All primers and sgRNA sequences used in this study are summarized in Supplemental Table S3e.

#### Plasmid construction and transgenic plant generation

For the generation of the *TurboID* (*TID*) and *WPRa4-TurboID* construct, the *TurboID* gene was cloned from the plasmid reported in the first *TurboID* study (ref. (26)) and tagged with 8x Histidine at the C-terminus. The gene cassette of *TID* and *WPRa4-TID* were then inserted into a binary vector pER10 and transformed into *Col-0* and *wpra4,* respectively, labeled as *TID*/*Col-0* and *WPRa4-TID*/*wpra4* in this study. T3 homozygous lines were selected using Kanamycin selection and combined in the proteomic analysis. These transgenic lines were applied for biotinylproteomic study.

For the generation of the transgenic plants that were subjected to the microscopic study, the full-length WPRa4 protein was fused with a yellow fluorescent protein (pIYFP, AY653732) (ref. (27)) at the N-terminus. The fusion gene was inserted into a binary vector (pHUB10) (ref. (28)) and placed under the control of a double cauliflower mosaic virus (CAMV) 35S promoter (d35SPro), and another construct driven by a native WPRa4 promoter as well. The two recombinant plasmids were subsequently transformed into the *wpra4* mutant to generate a group of T1 transgenic plants, d35SPro::His::YFP::WPRa4/*wpra4,* and WPRa4-Pro::His::YFP::WPRa4/*wpra4* respectively, as described in (12). These transgenic lines were applied for the subcellular localization study of WPRa4.

#### Classification of different plant force-responsive phases

The response to mechanical stress involves a series of morphological, physiological, and biochemical adaptations (29). We purposely divided the signaling events into four different phases according to data presented in the previously published papers: (1). Instant Force Response (IFR) is defined as the force response occurred within 1 minute of the force stress loading (in which Ca^2+^ spike and early phosphorylation signal are activated (11, 12)); (2). Prompt Force Response (PFR) is defined as the force response occurred between 1 and 5 minutes after the force stress loading (in which phosphorylation decayed while transcription started (12, 14)); (3). Rapid Force Response (RFR) is defined as the force response occurred between 5 minutes and 3.5 hours after the force stress loading (Touch responsive transcripts alteration and more than 90% of those transcripts peaked and returned to still state level (7, 13)); (4). Slow Force Response (SFR) is defined as a force response occurred from 3.5 hours to days after the force stress loading (7).

#### Antibody production

Rabbit polyclonal antibodies were raised against the S625-phosphorylated peptide of WPRa4(GL Biochem Ltd. Shanghai, China). The synthetic oligopeptide sequences of non-phosphorylated WPRa4 and the phosphorylated *wpra4* (pWPRa4), that were used to produce the polyclonal antibodies, was _619_VLMPNLSGIFNR and _619_VLMPNL***p*S**GIFNR, respectively. Rabbit polyclonal anti-PIP2A antibody was made as described before(30).

Anti-TurboID (AS20 4440) antibody was purchased from Agrisera, Anti-biotin (ab201341) and Anti-aequorin (ab9096) antibody was purchased from Abcam.

#### Confocal laser microscopy imaging of YFP::WPRa4 fusion protein

The YFP fluorescence and chlorophyll auto-fluorescence were examined in plant cells expressing His::YFP::WPRa4fusion protein under an Olympus IX81 laser scanning confocal microscope controlled by the Fluoview FV1000 system (Olympus America, Inc., Melville, NY, USA). The scan mode was XY while the auto-confocal aperture (C.A.) was opened. C.A. was 80 and 120 μm for the 20 x objective lens (0.75 NA) and 40 x objective lens (0.95 NA), respectively. For 60 x water immersion objective lens (1.2 NA), C.A. was 140 μm. YFP and chlorophyll auto-fluorescence were excited with a 515 nm and 635 nm argon laser line, respectively. The collecting emission ranges were 530 – 570 nm and 655 – 755 nm, respectively. Hypocotyl endodermal cells, leaf mesophyll cells, leaf guard cells, and root cells were obtained from one-week-old light-grown seedlings. Shoot apical meristem cells in flower buds from 5-week-old plants were examined.

#### Super-resolution microscopy imaging by dSTORM

During the dSTORM signal acquisition, excitation laser intensities (2.1 kW/cm^2^ and 4.5 kW/cm^2^ for 561 nm and 750 nm laser, respectively) were used to minimize photobleaching during imaging. Blinking signals from both channels were recorded with the EMCCD (100× EM gain, exposure time 30ms/ frame for 10,000 frames) simultaneously. Because the dye molecules briefly cycled from dark to bright and back to dark again for many iterations in the imaging buffer, this “winking” feature was used to calculate the two-dimensional (2D) Gaussian distribution. It was assumed that “winking” is centered at the location of a single dye molecule. Images were processed and analyzed using Rohdea 2.0 software (Nanobioimaging Ltd., China) and reconstructed with ImageJ software (National Institutes of Health, USA). All cell samples for dSTORM imaging were immobilized on Poly-L-Lysine coated coverslips and mounted in freshly prepared dSTORM imaging buffer: 50 mM Tris [pH 8.0], 10% [w/v] glucose, 560 μg/ml glucose oxidase, 57 μg/ml catalase, 2 mM cyclooctatetraene, 25 mM TCEP, 1 mM ascorbic acid, and 1 mM methyl viologen). The final resolution is determined to be < 20 nm in both channels based on an average fitting error.

#### Chloroplast isolation

The chloroplast isolation followed a published percoll gradient protocol(31). Briefly, 14-day-old wild-type (ecotype *Col-0*) light-grown seedlings were harvested and suspended in a chloroplast isolation buffer (0.6 M sorbitol, 10 mM MgCl_2,_ 10 mM EGTA, 10 mM EDTA, 20 mM NaHCO_3_, 40 mM HEPES pH 8.0) and homogenized in a cold room. The homogenate was subsequently filtered through a double-layer nylon cloth and centrifuged at 1,000 x g for 5 min to collect pellets. The resuspended pellet was applied on top of a pre-centrifuged continuous percoll gradient and centrifuged at 7,800 x g for 10 min at 4 ℃. Intact chloroplast layer was collected and washed with 3 x volumes of HMS (50 mM HEPES pH8.0, 3 mM MgSO_4_, 0.3 M sorbitol) buffer to remove excess percoll. The collected chloroplasts were subsequently centrifuged and resuspended in a small volume of HMS buffer for further protein extraction. The purity of isolated chloroplasts was examined under both an optical microscope (Nikon Eclipse TS100) and a confocal microscope (Zeiss LSM 710).

#### Biotinylated peptide sample enrichment and LC-MS/MS identification

For the TurboID-based PL experiment, homozygous transgenic plants of *TurboID*/*Col-0* and WPRa4*-TID*/*wpra4* were applied in this study. 21-23 days old plants grown on MS-Agar (Murashige and Skoog medium supplied with 10 g/L sucrose, and 8 g/L Agar) with 50 μM β-estradiol and 200 μM biotin were treated with a single-armed touch-machine for 20 min (Wang, K. 2018; Wang, K et al., 2019), then frozen with liquid N_2_ followed by protein extraction using Urea Extraction Buffer (UEB, with 150 mM Tris-Cl pH7.6, 8 M urea, 1.2 % Triton-X100, 2% SDS, 20 mM EDTA, 20 mM EGTA, 50 mM NaF, 1% G-2-P, 5 mM DTT, 1 mM PMSF, Complete EDTA free protease inhibitors cocktail, 5 mM ascorbic acid, and 2 % PVPP) (24, 32). The extracted protein samples were then precipitation with 12:1 (v/v) acetone: methanol cold solution at -20℃ overnight, centrifuged at 18,542 * g for 20 min, and then resuspended in the resuspension buffer (8 M urea, 50 mM Tris-HCl (pH 8.0), 0.2% sodium dodecanoate and 5 mM DTT), the precipitation procedure was done twice to remove free biotin as much as possible.

Protein was digested with trypsin (1:20 w/w) directly after extraction, followed by dimethyl labeling and respective sample mixing as described in (ref. (23)). Peptide samples from either *TurboID* or *WPRa4-TID* were diluted to 1mg/mL in 100 mM sodium acetate (pH 5.5) and then split into two fractions. One fraction was labeled with the ^12^CH_2_O (formaldehyde as the light labeling reagent), while the other fraction with ^13^CD_2_O (heavy isotope formaldehyde as the heavy labeling reagent). The ’Forward (F)’ was prepared by mixing light-labeled *TurboID* peptide and heavy-labeled *WPRa4-TID* peptide, and the switched combination was mixed as ‘Reciprocal (R)’. The labeling efficiency of dimethyl labeling is 99.6% for *Arabidopsis* peptides. Those mixed samples were then desalted and processed with tandem streptavidin enrichment. Briefly, 500 μL of beads was prewashed and balanced with Tris-buffered saline for 30 mins and transferred to the mixed samples for biotinylated peptides binding. The beads were then washed with TBS with 5% ACN for 5 times and ddH_2_O twice to remove the non-specific binding. The peptides were eluted with 0.2 % TFA, 0.1 % FA, with 80% ACN 5 times and combined. The high-purity biotinylated peptides sample was then desalted with Spin-tip (Thermo Scientific # 84850) and submitted for Orbitrap Eclipse Tribrid LC-MS/MS identification.

### LC-MS/MS analysis

The biotinylated peptides were reconstituted with solvent A (0.1% formic acid) and analyzed on an Easy nLC system (EASYSPRAY HPLC C18 COLUMN. 2UM, 100A, 75UM X 250MM) with a flow of 0.3 μ L/min. The biotinylated peptides were separated on a 60-minute gradient of 2 to 40 % solvent B (0.1% formic acid in 80% Acetonitrile): 2 to 30% solvent B, 0-53 minutes; 30% to 40% solvent B, 53-59 minutes; 40% solvent B to 100 % solvent B, 59-60 minutes. MS1 resolution was set to 60,000 with a scan range of 350–1500 m/z for a maximum injection time of 50 ms, while MS2 resolution was at 15,000 under HCD collision mode with normalized collision energy at 30% for a maximum injection time of 22 ms, the FAIMS mode was set to 2 CVs (-45V and -65V).

### Calculation of Biotin Occupancy of biotinylprotein

In this study, the biotinylated protein lysine occupancy was considered as a parameter to evaluate the accessibility of a protein to certain TurboID fusion proteins. The calculation equation is attached below. Calculated data were extracted from (Supplementary Table 1c). For either WPRa4-TID or TurboID labeled biotinylproteins, the <u>B</u>iotin <u>O</u>ccupancy (BO) is calculated as:

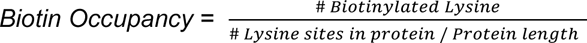

Thus, the ratio of (WPRa4-TID:TurboID) regarding the bProtein lysine Occupancy (Biotin Occupancy Ratio, BOR) is:

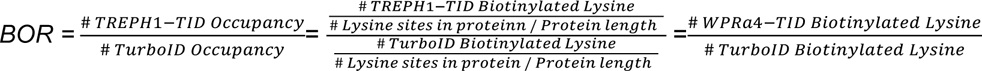

For these biotin occupancy values equal to 0 from either WPRa4-TID or TurboID, the corresponding occupancy values were set as half of the minimum (i.e. 0.0035 for WPRa4-TID and 0.0025 for TurboID) to facilitate the calculation.

### Quantitative biotinylproteomics

Raw data files (.raw) from Orbitrap Eclipse Tribrid Mass Spectrometer were converted to (.mgf) file using MS convert (Version: 3.0.20236-400e34473) for mascot Daemon (Version: 2.7.0, Matrix Science) search against TAIR_10 database and applied target/decoy strategy as reported previously in (Käll, L., et al., 2007). The searching parameters were: trypsin as protease, the peptide tolerance ± 10 ppm, MS/MS tolerance ± 0.02 Da, maximum missed cleavage set to 2, fixed modification included carbamidomethyl on cysteine, while variable modifications included oxidation on methionine, light dimethyl (H(4)C(2), 2x dimethyl = +28 Da) labeling was on lysine residue and N terminus of a peptide, heavy dimethyl (^2^H(4)^13^C(2), 2x dimethyl = +34 Da) labeling on lysine residue and N-terminus of a peptide, biotinylation (+226 Da) and biotin sulfoxide (+242 Da) on lysine residue. A percolator filter was applied and extracted ion chromatograms (XIC)-based protein level quantitation was performed in the in-house built software SQUA-D (Stable isotope-based Quantitation-Dimethyl labeling, Version 1.0; (ref. (23))). PSMs with q-value ≤ 0.01 after the percolator would be considered for a further quantification selection and the leading protein from the protein group was selected by default manner in the percolator. The selection criteria for quantifiable protein to be processed by SQUA-D software were the following: the PSM number of heavy dimethyl-labeled PTM peptides was ≥ 1, the PSM number of light dimethyl labeled PTM peptides PSMs ≥1, the number of different experimental replicates ≥ 4, the Mascot delta score ≥ 10 and the MS1 tolerance < 0.05 Da. Batch effect adjustment was applied to eliminate the variance caused by the experimental procedure in the quantification of 6 batches of samples (CF1, CF2, CF3, CR1, CR2, and CR3). The WPRa4 significantly proximal biotinylproteins (SPBs) were selected with the following criteria: Log_2_ ratio > 2.0, experimental replicates ≥ 5, biological replicate = 3, and the number of Log_2_ ratios of heavy and light dimethyl-labeled peptides ≥ 4, respectively. Here we use a highly stringent cut-off of Log_2_ ratio > 2.0 to keep the endogenous biotinylated protein eliminated. This cut-off is more stringent than most proximity-labeling proteomic studies. Those proteins that passed the selection criteria were defined as WPRa4-significantly proximal biotinylproteins (SPBs). Putative interactors were further defined by two criteria: first, quantitative Log_2_ ratio > 2.0, experimental replicates ≥ 5, biological replicates = 3 (SPBs criteria); second, Log_2_ (BOR) > 0.

### Mapping and clustering of putative interactors of WPRa4

The high-confidence WPRa4 putative interactor list was submitted to the STRING website ((33) accessed on Oct 2022) and the BIOGRID database (34) accessed on Oct 2022) and only those interactions with a confidence score greater than 0.7 from STRING were kept for further mapping, combined with interaction found in BIOGRID. NCBI homoloGene (https://www.ncbi.nlm.nih.gov/homologene/; build68, accessed on Oct 2022) database was also included to find certain genes with similarity. The network was then constituted and visualized in Cytoscape (Version 3.80). Interactor clustering was manually corrected and grouped with known gene function based on the Gene Ontology analysis using the GENE ONTOLOGY (http://geneontology.org/, accessed in Dec 2022), the subcellular localization was predicated by SUBA4 (Subcellular Localization Database for *Arabidopsis* Proteins 4 (35); accessed on Oct 2022).

For further plastid components and plastid movement components PPI map, PL result from this study in the “plastid related” group, combined with reported dataset XL-MS orthologues database (36) and BIOGRID (34) were included and visualized in Cytoscape (Version 3.80). PPI from different sources were marked accordingly for PL study (with inheriting node color and size from Figure 3D), recorded PPI contributor of other reported dataset XL-MS orthologues database (Dai et al., 2023) (Dash lines) and BIOGRID (parallel lines).

### Visualization of biotinylated sites on protein structures

The protein structure presented in this work comes from an online toolbox with ‘AlphaFold-Multimer’ service-based(37). Protein sequences were submitted to a public COSMIC² science gateway from (https://cosmic-cryoem.org/). For the coiled-coil protein interaction pattern prediction, WPRa4, WEB1, and PMI2 sequences from UniProt (https://www.uniprot.org/, accessed on Oct 2022) were subjected to the platform. The biotinylation statuses on four different types of structural domains, classified as non-loop (α-helix, β-sheet, turn) and loop regions in this study(38). Individual protein structures were downloaded from the AlphaFold database (39) (WPRa4, Q9LVQ4; WEB1, O48724; PMI1, Q9C8E6; PMI2, Q9C9N6). TIC-TOC complex was downloaded from RSCB PDB (PDBid: 7VCF) and the Tic100 protein from *Arabidopsis* was aligned with Chlamydomonas Tic100 to localize the biotinylated lysine position. Protein structures were visualized in Pymol 2.4.1 (Schrodinger, 2015). The secondary structure analysis is done with the DSSP algorithm(38) in R script (RStudio 2023.03.0+386 for Windows).

### Protein extraction and immunoblot assays

Protein samples prepared for immunoblotting were extracted from 17-day-old wild-type seedlings and the isolated chloroplasts using UEB, precipitated with 12:1 acetone: methanol, and resuspended in the resuspension buffer (RSB) (50 mM Tris–HCl (pH 6.8), 8 M urea, 1% SDS, 10 mM EDTA and 5 mM DTT). Extracted protein samples were loaded in equal amounts (30 μg per lane) side by side on 8% SDS-PAGE. The well-separated proteins were then transferred onto a PVDF membrane (Millipore). Anti-WPRa4 and Anti-actin antibodies were incubated and detected as previously described in (ref.(12)). Relative protein abundance was measured using ImageJ software.

### Aequorin and Ca^2+^ spike-monitoring

Crispr knock-out homozygous mutants against *AEQ* expression plant background were used for Ca^2+^ spike signal-monitoring. About 20 seedlings grown on agar together were added 30 μL 5 mM Coelenterazine-h (Cat.C6780, Thermo, USA) for overnight before imaging. A 40-touch treatment was applied to the tested plant within one minute (defined as the IFR phase) using an air-pump-driven cotton touch system in the monitoring dark box. The temporal profile of luminescence by cotton touch was detected and recorded through a custom-built photon-multiplier tube (PMT) platform including a P10PC PMT (Electron Tube Enterprises, Uxbridge, UK) which was mounted in a dark box (provided by Science Wares, East Falmouth, MA, USA). *AEQ/Col-0*, *AEQ*/*pmi4-c*, *AEQ*/*arc3-c*, *AEQ*/*arc3pmi4-c* and *AEQ*/*piezo1-c plants* were applied to test the touch-induced Ca^2+^ oscillation.

### Automated Hair Touch treatment

Thigmomorphogenesis (a Slow Force Response of a plant, SFR) under mechanical stimulation for days were examined in *Col-0*, *arc3, pmi4,* and *arc3pmi4* by an automatic hair touch-force loading machine (17). The 14-day-old plants grown in soil under constant light conditions (150 - 260 µEm^-2^s^-1^) were subjected to long-term (> 12 days, depending on the bolting time of the tested plant) mechano-stimulation *i.e.* 4 rounds of touches a day. In each round, the human hair brushes touched plants 40 times within 20 rounds of trips and applied 1 - 2 mN force. One round of touch treatment lasted for 3 minutes. The size of the plant was record one day before treatment and six days after treatment by phone and analyzed by ImageJ (version 1.53c). When the height of the primary inflorescence reached 1 cm, the plant was recorded as a bolted plant. Long-term mechanical stimulation lasted 20-22 days for four biological experiments. *Col-0* and the T-DNA insertion mutants *pmi4*, *arc3*, and *arc3pmi4* were used in the automated hair touch assay.

### Transcriptomics and RT-qPCR quantification of gene expression

In the transcriptomic analysis of *pmi4* and *Col-0* at both PFR and RFR phases, samples were harvested on the early PFR phase (touched for 3 min using the automated hair touch) and early RFR phase (touched for 3 min using the automated hair touch and waiting for 17 min) between *Col-0* and *pmi4*in three biological replicates. Trizol buffer was applied to extract total RNA for sample preparation of transcriptome analysis. The 100 μL of fine ground tissue powder was dissolved in 1 mL Trizol buffer and mixed under vortex for 10 -15 sec. The mixing buffer was dissociated at room temperature for 10 min. 200 μL chloroform was added to the dissociation buffer under vortex for 15 sec violently and waited for 10 min. The buffer would be separated into two parts by centrifugation of 11,000 g for 15 min at 4 °C. 500 μL transparent supernatant was transferred into a new tube and mixed gently with 500 μL isopropanol as well as 100 μL NaAC (Sodium acetate, PH5.2). The mixing buffer was kept at 4 °C for at least 30 min. Through centrifugation of 11,000 g for 10 min at 4 °C, RNA was precipitated on the bottom of the tube and 1 mL 70% ethanol was used to wash the RNA. After the washing step was repeated once, ethanol was removed, and RNA was dried for 5-10 min. 50 μL DEPC treated water was added to dissolve RNA and 50 μL isopropanol as well as 10 μL NaAC was mixed with the RNA to precipitate again. Through washing and drying step, the RNA was dissolved in 20-30 μL DEPC treated water waiting for measuring the concentration.

The concentration of the RNA was measured by NanoDrop 2000 (Thermo Scientific, Waltham, USA). 5 μg RNA was dilution with 5 μL10X DNase I Reaction Buffer into a 50 μL reaction system. 1 μL (2 units/μL) DNase I was mixed with the reaction system and incubated at 37 °C for 15 minutes. 1 μL 0.5 M EDTA was added into the buffer and heated at 75 °C for 10 minutes to inactivate the enzyme. The RNA integrity was confirmed by both formaldehyde agarose gel electrophoresis and 5200 Fragment Analyzer (Agilent Technologies, CA, USA).

The cDNA library was prepared and sequenced by BGI company (China) through the DNBSEQ platform. Low-quality, adaptor-polluted, and high content of unknown base (N) reads were removed before downstream analyses. The clean reads were mapped to the reference genome (*Arabidopsis*_thaliana, GCF_000001735.4_TAIR10.1) using Bowtie2 through HISAT method (40). Gene expression analysis was conducted by RSEM (41).

In the RT-qPCR experiment, the sample preparation was carried out based on standard protocols. Tissue samples from *Col-0* and *pmi4*were harvested at different time point (5 min as in PFR, 15 min as in RFR, 5 h and 10 h as in SFR). The extracted RNA was incubated with DNase I (New England Biolabs, Massachusetts, USA) to remove DNA contamination and reversely transcribed into cDNA using a SuperScript III First-Strand Kit (Invitrogen, Thermo Fisher Scientific, USA). The quantitative PCR was performed on a LightCycler® 480 instrument II (Roche, Basel Switzerland) through LightCycler 480 SYBR Green I Master (Roche) based on a 20 μl reaction volume system with a 384-well plate according to the manufacturer’s specifications. The PCR program included pre-incubation at 95 °C for 10 min, 45 cycles of denaturation at 95 °C for 10 s and annealing at 60 °C for 15 s, and last extension at 72 °C for 18 s. Primer3 (https://bioinfo.ut.ee/primer3/) and NCBI primer design (https://www.ncbi.nlm.nih.gov/tools/primer-blast/) were used to design gene-specific primers (22, 42). The cycle number at the threshold was applied for gene expression quantification based on the previous 2-ΔΔCt method (43).

*Col-0* and the T-DNA insertion *pmi4* mutant were used for transcriptome analysis. The primers were summarized in Supplemental Table S3e.

## Results

### Microscopic and biochemical studies revealed that WPRa4 is associated with plastid

Two well-characterized cytosolic WPR family proteins that contain a central DUF827 domain composed of a long coiled-coil region for protein-protein interaction, WEB1, and PMI2, have been reported to be involved in plastid movement (44). However, the cytoskeletal WPRa4 protein has not been demonstrated to be associated with plastid (16), but characterized as a touch-responsive phosphorylated protein involved in plant mechanoresponse (12). To further confirm the subcellular and cytosolic localization of WPRa4, we examined the bioluminescent signal emitted from the native promoter-driven fusion protein YFP-WPRa4 using confocal microscopy (Fig.1A). In mesophyll cells, it was evident that the YFP-WPRa4 was localized both on chloroplasts and plasma membrane (Fig.1A). The YFP-WPRa4 was also visualized in the cytoplasmic region periphery to the plasma membrane of epidermal cells, hypocotyl cells, and root elongation zone cells (Supplemental Fig. S1). Similar localization was also observed under double 35S promoter-driven YFP-WPRa4 transgenic plants (Supplemental Fig. S2).

**Figure 1.**
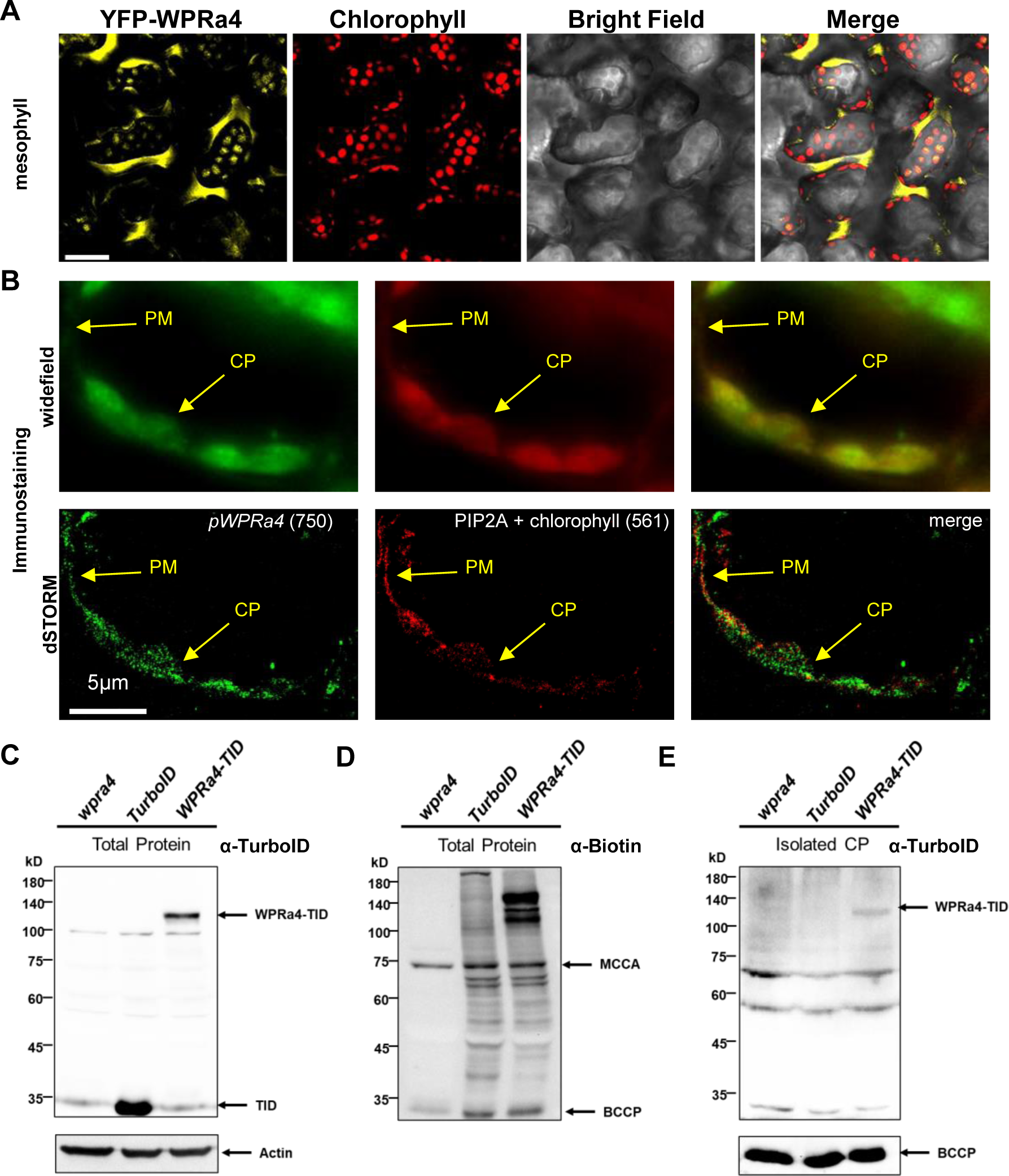
WPRa4 protein is associated with chloroplast. (A) Subcellular distribution of WPRa4 in *Pro::His::YFP::*WPRa4*/wpra4* transgenic plant mesophyll cells. Scale bars are 30 μm. The excitation of YFP and chlorophyll auto-fluorescence were performed with 515 nm and 635 nm argon lasers, respectively. More confocal images can be found in Supplemental Fig. S1 and Fig. S2. (B) Super-resolution microscopy examination of the subcellular localization of pWPRa4 and PIP2A proteins in leaf cells. From the left panel to the right, fluorescence is from *p*WPRa4 (synthetic green), PIP2A + chlorophyll auto-fluorescence (synthetic red), and co-localized merged photograph. The 750 nm laser channel was used to observe *p*WPRa4, while the 561 nm laser channel was used to observe PIP2A. PM, plasma membrane; CP, chloroplast. (C) Immunoblots showing the TurboID fusion protein expression pattern of 17 days seedlings *wpra4, TurboID/Col-0, and* WPRa4*-TID/wpra4*. Anti-TurboID antibody (Agrisera AS204440) was used as the primary antibody. WPRa4-TID band (∼120 kD) and TurboID band (∼35 kD) were marked accordingly. The letter α stands for ‘anti’. (D) Immunoblots showing the biotinylated protein pattern of 17 days seedlings

We further performed the dual-color dSTORM super-resolution microscopic imaging of both WPRa4 protein and water channel proteins in mesophyll cells using both the phosphorylated isoform, WPRa4-pS625 (phosphorylation on serine 625), -specific polyclonal antibodies (12) and the PIP2A (plasma membrane intrinsic protein 2A, an plasma membrane aquaporin protein) protein-specific polyclonal antibodies (30). The luminescent light signals from anti-S625 site-phosphorylated WPRa4 antibodies were clustered well with those antibodies against the water channel PIP2A, while the co-localized luminescent light signals with the chloroplasts were also observed but of a less dominant intensity (Fig.1B).

To further confirm the chloroplast localization of WPRa4 and to characterize the proximal proteins of WPRa4, we also isolated chloroplasts from each of both *TurboID (TID)*/*Col-0* and WPRa4*-TID*/*wpra4* transgenic plants (Fig.1C, 1D). Using an anti-TurboID polyclonal antibody and the untransformed *wpra4* mutant plant as the controls, the immunoblots confirmed the presence of WPRa4-TID fusion protein in the highly isolated chloroplast samples, while the TurboID protein was absent (Fig. 1E, Supplemental Fig. S3). The three experiments described provide evidence that the cytoskeletal protein WPRa4 is associated with plastids in Arabidopsis, suggesting a potential network, which involves a WPRa4-harboring continuum comprising of the cytoskeleton and plastoskeleton, may function in mechanotransduction.

### Quantitative biotinylproteomics study found that WPRa4 might interact with numerous plastid-related cytoskeletal proteins

As WPRa4 possesses neither a transmembrane domain nor a plastid transit peptide sequence, we speculated that the special plastid localization of WPRa4 is probably through protein-protein interaction with a plastid envelope-localized protein. The proximal protein network composition of WPRa4 was therefore determined using a combinatorial dual-labeling procedure, consisting of both *in planta* <u>p</u>roximity labeling (PL) and *in vitro* dimethyl labeling, enabling quantitative biotinylproteomic analysis (See Methods and Supplementary Materials).

A graphic workflow of this approach was presented in Supplemental Fig. S4. In short, three biological replicates of peptide preparations performed on the *TurboID/Col-0* and *WPRa4-TID/wpra4* plants (Fig. 2A), which were grown on Murashige and Skoog agar medium supplied with 200 μM biotin, yielded a total of 6 mass spectrometer injection samples with high biotinylpeptide purity (Supplemental Fig. S5A-B, Supplemental Table S1-0). All 16,266 PSMs acquired from 6 injections were presented in Supplemental Table S1a, and 13,846 biotinylated PSMs were selected according to previously published standards (45) and presented in Supplemental Table S1b for further analysis. As a result, 2136 label-independent (labeled with both light and heavy formaldehyde) and repeatable (PSM ≥ 2) biotinylated peptides (Supplemental Table S1c), derived from 1267 biotinylproteins, were identified (Fig. 2B, 2C, Supplemental Table S1d). Among those 1225 TurboID labeled proteins and 763 WPRa4-TID labeled proteins (Fig. 2C, Supplemental Table S1e), 27.1 % and 23.6 % were biotinylated on multiple lysine sites, respectively (Fig. 2D, 2E, Supplemental Fig. 5C). Notably, the most extensively labeled biotinylprotein by the WPRa4-TID fusion biotin ligase was WPRa4 itself (30 biotinylated sites, Fig. 2F, 2G). The PSMs (peptide-spectrum matches) of each biotinylated peptide distribution followed Zipf’s law, with a slope (k) of -0.33887 and -0.5065 for TID and WPRa4-TID, respectively (Supplemental Fig. 5D). Peptide length distribution analysis showed that most of the peptides attributed to 7 to 24 amino acids long for both TurboID (95.2 %) and WPRa4-TID (95.5%) (Supplemental Fig. 5E) labeled samples.

**Figure 2.**
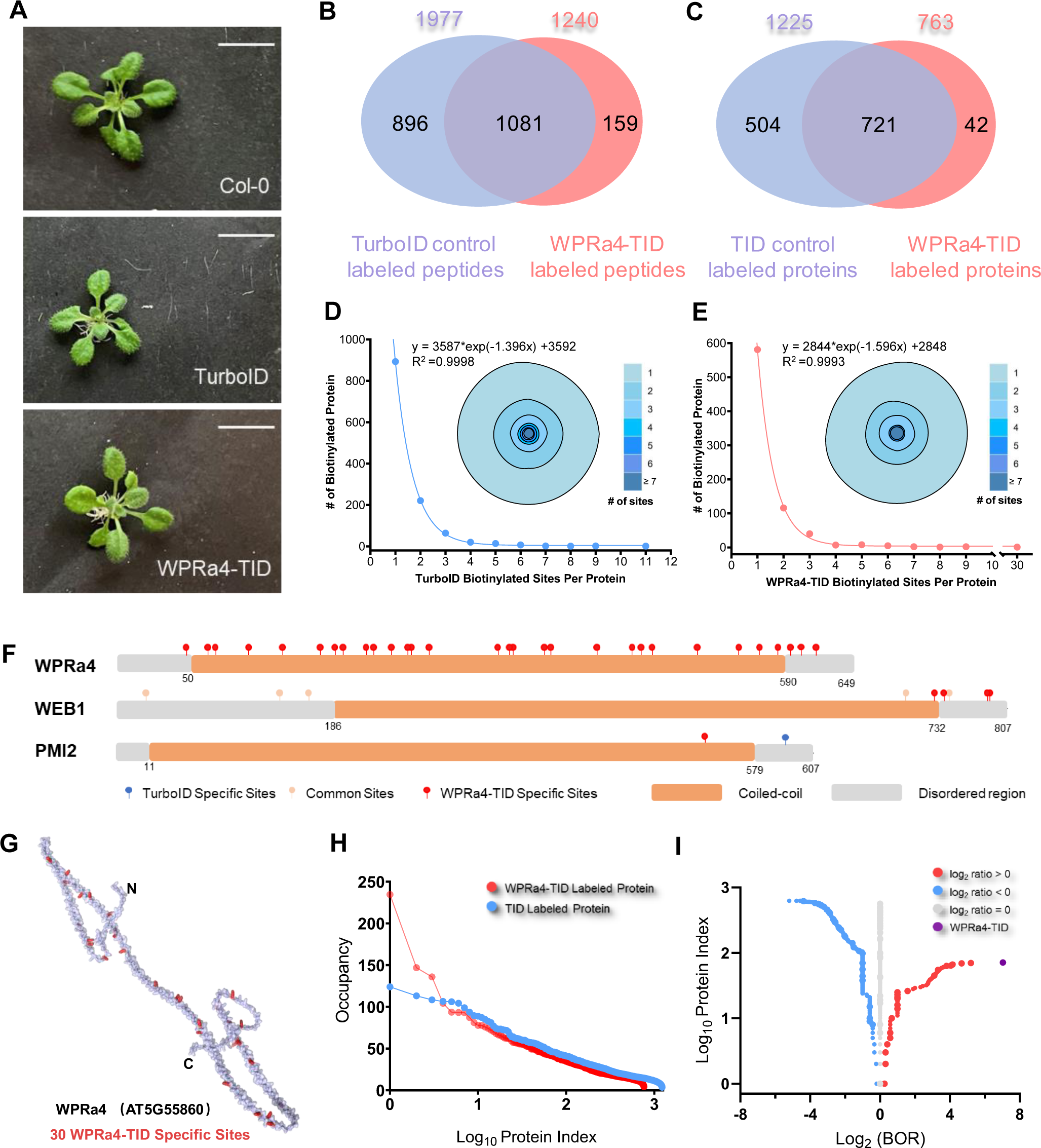
Biotinylproteomics analysis revealed several WPR proteins as proximal proteins of WPRa4. (A) Image of 24-day-old *TurboID* and *WPRa4-TID* plants grown on MS medium supplied with β-estradiol and biotin, scale bar = 1 cm. (B) A Venn diagram shows the differential biotinylated proteins from *WPRa4-TID* transgenic plant (763) and *TurboID* transgenic plant (1225). (Original data can be found in Supplemental Table S1e). (C) A Venn diagram shows the 2136 biotinylated peptides from the *WPRa4-TID* transgenic plant (1240) and *TurboID* transgenic plant (1977). (D-E) Curve and tree ring showing the distribution of TurboID (D) and WPRa4-TID (E) labeled protein by biotinylated site number. (Original data can be found in Supplemental Table S1d). The data is fitted with an exponential decay equation model with non-linear regression. A merged plot is shown in Supplemental Fig. S5C. (F) A diagram shows the biotinylated sites on three representative proteins, WPRa4, WEB1, and PMI2. Coiled-coil sequence features were referred from AlphaFold predicted structure (WPRa4, Q9LVQ4; WEB1, O48724; PMI2, Q9C9N6). Sites distribution on the 3D structure of WEB1 and PMI2 can be found in Supplemental Fig. S5G. (G) Displaying of biotinylated sites on the AlphaFold predicted WPRa4 structure, red residuals indicate the biotinylated lysine sites. (H) Distribution of the biotin occupancy (BO) of biotinylated proteins labeled by TurboID (blue) and WPRa4-TID (red). (I) A volcano plot showing the biotin occupancy ratio (BOR) distribution with protein index from biotinylprotein identified in this study. The protein index is ranked by BOR value for the separated dataset of Log_2_(BOR) greater than 0, equal to 0, and less than 0. (Original data can be found in Supplemental Table S1f).

Topological analysis on those TID-labeled and WPRa4-TID-labeled biotinylated sites on several relevant proteins was highlighted with the AlphaFold (39) predicted protein structures. Consistent with previously published topological analysis results (22), more than half of biotinylated sites were localized on the loop region from TID-labeled (980 sites, 55.4%) and WPRa4-TID-labeled biotinylproteins (586 sites, 53.3%) (Supplemental Fig. 5F). Two previously reported WPR proteins, WEB1and PMI2, were differentially labeled by both TurboID biotin ligase and WPRa4-TID fusion biotin ligase (Fig. 2F, Supplemental Fig. 5G). PMI1 (Plastid Movement Impaired 1) was also identified with 4 differentially labeled biotinylated sites (Supplemental Fig. 5H). Visualization of plastid movement related WEB1, PMI1, and PMI2 proteins using the biotinylated site information revealed 11 biotinylated sites on the loop regions while only 4 sites were found on the non-loop regions, which supports a loop structure preference for the biotinylation activities of the promiscuous biotin ligase (Supplemental Fig. 5H).

Using the lysine percentage-dependent Biotin Occupancy (BO) calculation method (22), these biotinylproteins identified from both TurboID- and WPRa4-TID fusion protein-harboring transgenic plants were analyzed subsequently. The results showed that the WPRa4-TID fusion ligase produced more biotinylproteins of higher Biotin Occupancy (BO) than those from TurboID transgenic plants although a lesser number of biotinylated peptides and proteins were identified from the *WPRa4-TID* transgenics (Fig. 2H). The Biotin Occupancy Ratio (BOR) (22) was also adopted to analyze biotinylproteins found from both TurboID and WPRa4-TID biotin ligase expression transgenics (Fig. 2G, Supplemental Table S1f) in order to determine the proximal biotinylproteins of T TurboID ID biotin ligase (Log_2_(BOR) < 0, 627 proteins), common biotinylproteins (Log_2_(BOR) = 0, 569 proteins), and the proximal biotinylproteins of WPRa4-TID fusion ligase (Log_2_(BOR) > 0, 71 proteins). The biotinylprotein of the largest Log_2_(BOR) of 7.04 was the WPRa4-TID fusion protein itself with 30 biotin-labeling sites on it according to the biotinylproteins identified from *WPRa4-TID* transgenic plants. This BOR parameter (Log_2_ (BOR) > 0) is thus considered one of the criteria for determining the extent of the proximity of biotinylproteins toward the bait protein WPRa4-TID.

An in-house developed quantitative PTM proteomics workflow (46) was applied in the analysis of 618 quantifiable biotinylproteins. They were specially selected according to a series of previously stipulated criteria including the number of peptide spectral match (PSM) (Supplementary Table S2a, and Materials and Methods). Furthermore, multiple hypothesis correction and the Log_2_ ratio cut-off (BH-FDR ≤ 0.05, Log_2_ ratio > 2.0, biological replicates = 3, experimental replicates ≥ 5) were applied on the Mass Spectrometry (MS) Extracted Ion Chromatography (XIC) data collected from both *TurboID* and *WPRa4-TID* transgenics when using the proteome quantitation software SQUA-D (23). As a result, 49 proteins were identified to be significantly proximal to the bait protein WPRa4-TID. They were defined as WPRa4-significantly proximal biotinylproteins (SPBs) (Fig. 2H, Supplemental Table S2a). In contrast, 46 biotinylproteins were significantly proximal to TID and were defined as TurboID SPBs (BH-FDR ≤ 0.05, Log_2_ WPRa4-TID/TurboID ratio < -2.0, biological replicates = 3, experimental replicates ≥ 5).

The Biotin Occupancy Ratios (BOR) and GO analysis results of these WPRa4-TID significantly proximal biotinylproteins were compared with those of TID, respectively, and a distinct trend was found between these two biotinylprotein lists (Fig. 3A, Supplemental Fig. S6A-B). Subsequently, we integrated the BO ratio (BOR) with the Log_2_ ratio of XIC ion intensity for each biotinylprotein to determine the putative interactors of WPRa4 based on two-dimensional comparison and standard selection (Fig. 3B-3C). As a result, 20 putative interactors of WPRa4 of both XIC ion intensity Log_2_ ratio > 2.0 and Log_2_(BOR) > 0 were selected (Fig. 3A-3C, Supplemental Fig. S6C, Supplemental Table S2b). Among these biotinylproteins, two kinases were identified from this study as putative interactors of WPRa4, CBC1 (Convergence of blue light and CO_2_, AT3G01490), and a novel receptor-like kinase named WR-RLK 1 (WPRa4-Related Receptor-Like Kinase 1, AT1G14390). It is notable that the WR-RLK1 has the largest XIC Log_2_ ratio among all those putative interactors of WPRa4.

**Figure 3.**
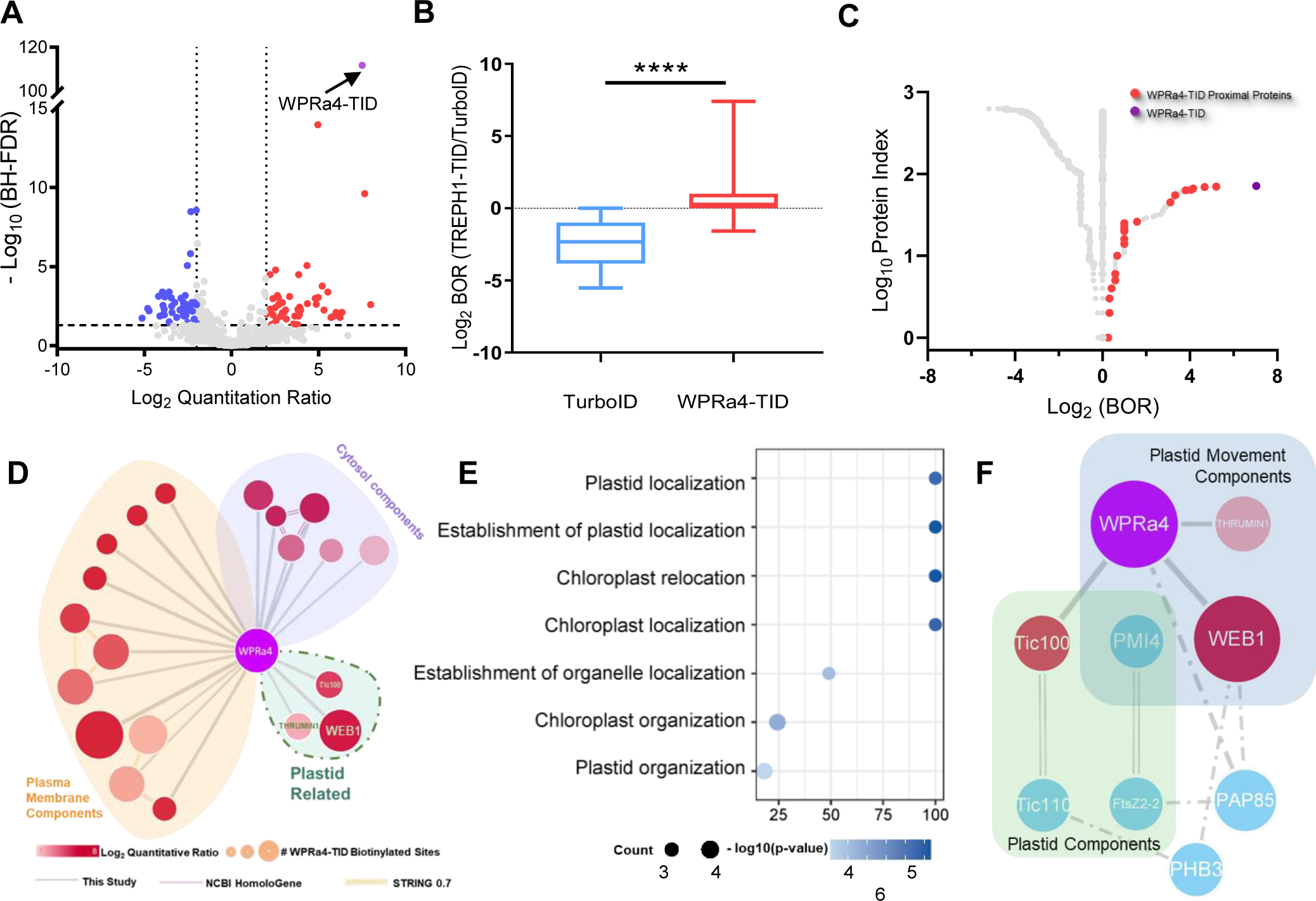
Quantitative analysis and bioinformatics analysis revealed that putative interactors of WPRa4 are enriched in plastids. (A) A volcano plot showing quantitative biotinylproteomic result of the WPRa4-TID and TurboID control. Proteins passed the selection criteria (BH-FDR ≤ 0.05, WPRa4-TID/TurboID log_2_ ratio > 2.0, biological replicates = 3, experimental replicates ≥ 4) were considered as WPRa4 proximal proteins (Red dots). (Original data can be found in Supplemental Table S2a). (B) A box plot showing the BOR distribution of TurboID proximal proteins (blue) and WPRa4-TID proximal proteins (red). Student t-test was applied to compare the difference. **** indicates the p-value < 0.0001. (C) A volcano plot showing the WPRa4 proximal proteins (red) and WPRa4-TID (purple) with a quantitative ratio > 2.0 and BOR > 0. The protein index is ranked by BOR value for the separated dataset of Log_2_(BOR) . (Original data can be found in Supplemental Table S2b). (D) Map of the 21 significant proximal proteins using Cytoscape. Line type indicates the source of connection from this study (grey), NCBI homologene database (purple), and STRING (yellow, with confidence over 0.7). The size of each protein indicates the Log_2_ Quantitative ratio. (Original data can be found in Supplemental Table S2b). (E) GO analysis on biological_process (BP) result using the 21 significant proximal proteins, with the x-axis indicating enrichment fold, y-axis indicating different terms, node size indicating the number of protein enriched, and color gradient indicating FDR corrected p-value. (F). Protein-Protein interaction map generated from plastid related group of Figure 3D. WPRa4, WEB1, THRUMIN1, and Tic100 were identified in this PL study (with inheriting node color and size from Figure 3D), and other blue nodes represent recorded PPI contributors of other reported dataset XL-MS (crosslinking mass spectrometry) orthologues database (Dai et al., 2023) (Dash lines) and BIOGRID (parallel lines).

### Bioinformatic analysis suggested a role for the plastoskeletal protein PMI4 in Arabidopsis mechanoresponse

To decipher the subcellular localization of WPRa4 based on the analysis of its putative interactors, a WPRa4-centered putative interactome was generated and further categorized into three subgroups (Fig. 3D): plasma membrane components (11 proteins), cytosol components (6 proteins), and plastid-related proteins (3 proteins) (35). Gene Ontology (GO) analysis of these 20 putative interactors of WPRa4 revealed that several were related to plastid localization and plastid organization, including WEB1, THRUMIN1 (Light-Regulated Actin-Bundling Protein Involved in Chloroplast Motility, AT1G64500), and Tic100 (∼100 kD plastid translocon subunit at the inner envelope membrane, AT5G22640) (Fig. 3E). We also noticed that Tic100 protein of plastid translocon TOC-TIC super-complex was specifically biotin-labeled by WPRa4-TID on K528 and K574 sites (Supplemental Fig. S6D), which localized in the intermembrane space of previous determined TOC-TIC supercomplex (47). These findings indicate that WPRa4 is localized in a position proximal to both the plasma membrane and the plastid. This conclusion is consistent with the observation conducted by microscopy (Fig. 1). Therefore, the quantitative biotinyproteomics data support a model, in which WPRa4 acts as a molecular connector, linking the cytoskeleton to the plastid through plastid envelope-localized proteins TOC-TIC complex, which is well-known to connect with the plastoskeleton, and thereby integrating the cytoplasmic skeletal proteins with plasma membrane and plastids to form mechanoresponsive signaling pathways.

Indeed, WPRa4 has been shown to be a mechanoresponsive signaling component, known as Touch Regulated Phosphoprotein 1 (12). Bioinformatic analysis also indicated the plastid-related WEB1, THRUMIN1, and Tic100 proteins (Fig. 3D) are putative interactors (or called proximal proteins) of WPRa4, which suggests that these plastid-related cytoskeletal proteins might work together or even with plastids to play the role in touch response. In addition, since a previously performed genetic screen for the defective plastid movement phenotype (19) has provided both the cytosolic and the plastidic *<u>p</u>lastid <u>m</u>ovement-impaired* (*pmi*) mutant candidates for investigating the roles of these mutants in plant touch response, we therefore analyzed possible protein-protein interaction (PPI) network from the public PPI database BIOGRID (34) and previously published orthologues XL-MS (crosslinking mass spectrometry) dataset (36). This bioinformatic analysis revealed two major protein groups: the plastid movement components group and the plastid component group, with the intersection protein being PMI4 (Fig. 3F). PMI4 is a previous *PMI* gene that encodes a plastoskeleton tubulin-like protein tethered with the chloroplast inner membrane and it interacts with FtsZ2 protein to form a Z-ring complex during chloroplast division (21, 48). FtsZ2 protein happens to be located both on the plastid envelope and in the stroma region where it interacts with the PMI4 protein (49).

Based on the results presented, there are four compelling reasons to investigate PMI4’s role in touch response through the bioinformatics analysis of biotinylproteomic and XL-MS-based interactomic results: (1) Biotinylproteomics shows that the cytoskeletal protein WPRa4 is located close to plastids, and several putative interactors are linked to plastid movement (Fig 2D, 2E). (2) Previous Genetic screening of mutants on plastid movement reveals that both cytosolic and plastidic skeleton proteins, including WEB1, THRUMIN1, PMI1, PMI2, and PMI4, are involved in this plastid movement processes (16, 19, 50). Interestingly, PMI4 is the only one of this kind of proteins that is localized within the plastid stroma and required for normal plastid movement. (3) These structural proteins related to plastid movement are connected to plastoskeletal proteins through XL-MS proteomics and bioinformatics analysis (Fig. 3F), where PMI4 is the intersection of the plastid component and the plastid motion component. (4) Organelles are known to play a role in mechano-sensing, with force signals potentially transmitted through cytoskeletal and plastoskeletal proteins (5) into subsequent signaling events.

Since the involvement of the cytoskeletal protein WPRa4 in touch response has been validated (12), we hereby focus on testing PMI4, a plastoskeletal protein, on its role in mechanosensing.

### PMI4 plays a crucial role in the plant’s touch response, influencing both touch-induced calcium oscillations and the touch-responsive transcriptome

The *pmi4* mutant has been shown to exhibit chloroplasts of enlarged size and reduced number in the mesophyll cells (20). As a control for the *pmi4* mutation-resulted phenotypes of both plastid size enlargement and plastid number reduction, another plastid division mutant, *accumulation and replication of chloroplasts 3 (arc3),* was selected as a control because of its similar phenotypes in the enlarged chloroplast size, the reduced number of plastids per cell and the multiple misplaced Z-rings, while the *PMI4* expression level found in *arc3* mutant is not affected (21). It is therefore that these two types of plastid mutants were used together to investigate the role of PMI4 protein in thigmomorphogenesis. The plastid morphologies of the T-DNA insertion mutants, *arc3, pmi4,* and *arc3pmi4* double mutant (Genotyping information was shown in Supplemental Fig. S7A-B), were analyzed together with that of the wild-type (*Col-0*) plant (Supplemental Fig. 7C, 7D, 7E, 7F) using confocal microscopy. The analysis revealed that the chloroplasts in the *arc3*, *pmi4*, and *arc3pmi4* double mutants exhibited an increased size, accompanied by a reduced number of plastids in these genotypes (Supplemental Fig. 7G, 7H).

Subsequently, these mutants were subjected to the automated hair touch treatment (12). Thigmomorphogenesis has been considered to be a <u>S</u>low <u>F</u>orce <u>R</u>esponse (SFR, see Material and Methods, “Classification of different plant force-responsive phases”). These results indicated that both *Col-0* and *arc3* exhibited a normal touch-induced reduction in rosette size, while both *pmi4* and *arc3pmi4* double mutant had an insignificant rosette size difference between the touched and untouched plants (Fig. 4A-D). At the same time, the *pmi4* mutant displayed an insensitive phenotype on the bolting delay (29.8 ± 0.1 d vs. 30.0 ± 0.1 d, Fig. 4g, Supplemental Fig. S10) as compared to that of the wild-type (*Col-0*) plant (29.8 ± 0.2 d vs. 30.8 ± 0.2 d, Fig. 4E, Supplemental Fig. S8). The *arc3pmi4* double mutant also showed a touch-insensitive phenotype (28.4 ± 0.2 d vs. 28.8 ± 0.2 d, Fig. 4H, Supplemental Fig. S11) like did the *pmi4* mutant. The *arc3* mutant alone exhibited a touch-sensitive phenotype, *i.e*. the reduced rosette size and delayed bolting (28.8 ± 0.2 d vs. 29.9 ± 0.2 d, Fig. 4F, Supplemental Fig. S9). Based on these observations, we concluded that the tubulin-like PMI4 protein rather than the altered plastid phenotypes played a crucial role in plant thigmomorphogenesis (*i.e.* bolting time and rosette size).

**Figure 4.**
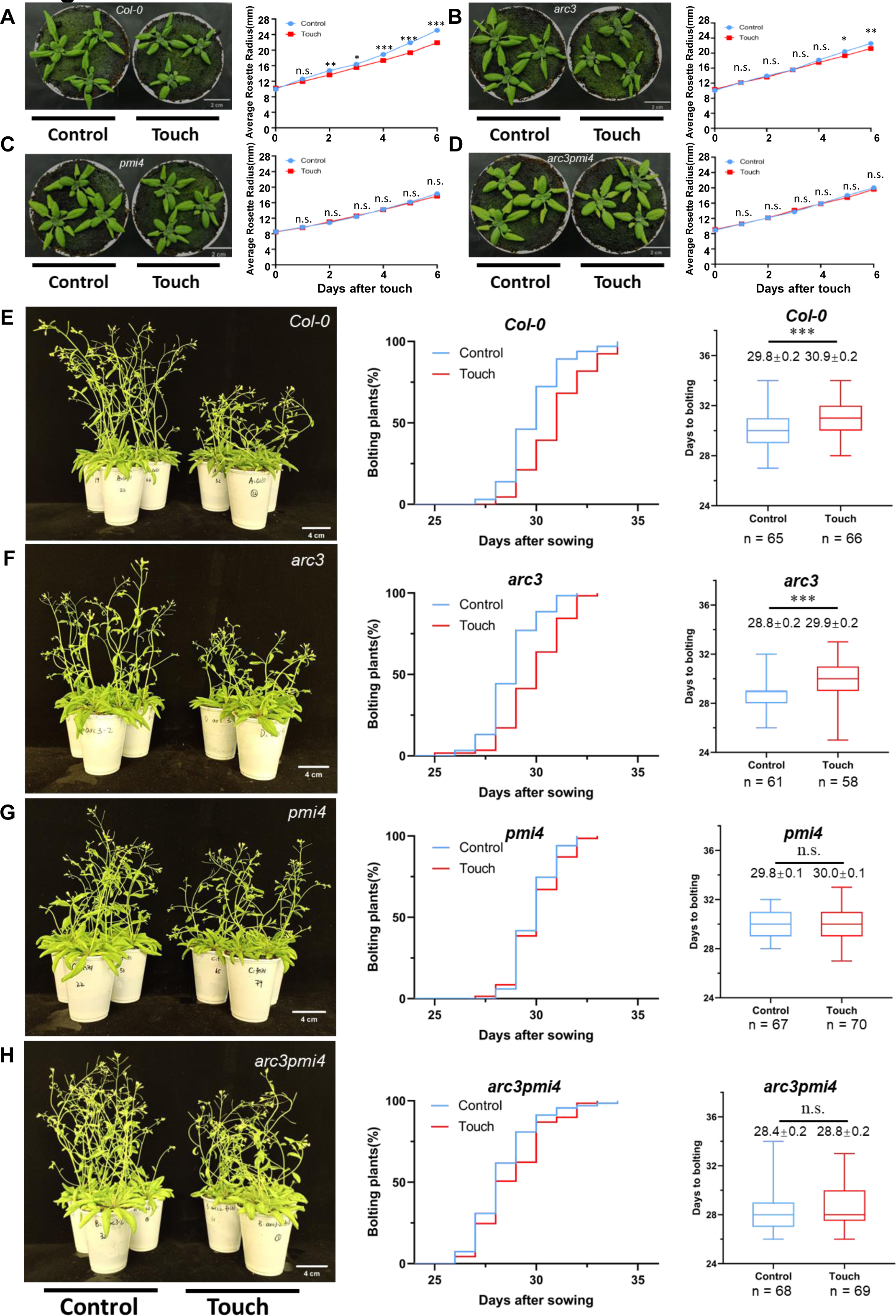
Touch response on rosette size reduction and bolting delay of selected chloroplast mutants. (A-D) Photos showing control (left) and touch (right) plant rosette differences in *Col-0* (A) and T-DNA insertion mutants of *arc3* (B)*, pmi4* (C), and *arc3pmi4* (D) plants. A comparison of the plant rosette size under control (blue line) and touch (red line) was shown on the right panel next to the photos. (E-H) Touch response of *Col-0* (E)*, arc3* (F)*, pmi4* (G), and *arc3pmi4* (H), with photos showing representative plants (left panel), Kaplan-Meier plots showing growth period (middle panel), Box-and-whisker plots showing the bolting date difference (right panel). Statistical significance between control and touched group plant for touch response was obtained using student t-test: (***, P values less than 0.001; **, P values less than 0.01; *, P values less than 0.05; n.s., not significant). Relating supplemental figures can be found in Supplemental Fig. S9-S12.

Touch is well-known to induce the cytoplasmic calcium spike (11), which is treated as an Instant <u>F</u>orce <u>R</u>esponse (IFR, the response occurs ≤ 1 min following stimulation). In the present study, as the aequorin (AEQ) is able to report the real-time Ca^2+^ concentration oscillation upon the touching treatment (11), we measured the calcium spike alterations in four transgenic plants, *AEQ/Col-0*, *AEQ*/*arc3-c*, *AEQ*/*pmi4-c*, and *AEQ*/*arc3pmi4-c* (Supplemental Fig. S12) to address the question if PMI4 protein influences the mechano-stimulated calcium oscillation. All three types of null mutants were generated using the CRISPR approach in an *AEQ* reporter gene over-expression background (See the Materials and Methods for detailed genotype information).

The treatment of 40 touches (See Materials and Methods for details) was subsequently applied to these mutants with the *AEQ* reporter gene. The chemiluminescence signals of the corresponding cytoplasmic calcium transient changes were captured and measured using a photonic reporting system (Fig. 5A; see Materials and Methods for details). The aequorin protein expression levels in these transgenic mutant lines were measured by immunoblots (Fig. 5B). Consistent with the bolting delay phenotype of *pmi4* mutants, the Ca^2+^ spikes were impaired with the touching in both *AEQ*/*pmi4-c* and *AEQ*/*arc3pmi4-c* transgenic backgrounds as compared to those of *AEQ*/*arc3-c* and *AEQ*/*Col-0* control transgenics (Fig. 5C, 5D), suggesting that *PMI4* gene alone plays a crucial role in regulation of the touch-induced Ca^2+^ transient, or the Instant Force Response (IFR). In contrast, *AEQ*/*arc3-c* mutant transgenics failed to suppress the Ca^2+^ spike as failed the *wpra4* mutant (12), indicating that this impaired Ca^2+^ spike is not caused by altered plastid size and number, but by the defects in the tubulin-like PMI4 protein inside the plastid. To eliminate the possible influence inflicted by the aequorin expression level, the signal peaks were normalized with aequorin expression levels in transgenic plants (Fig. 5E), indicating the impaired Ca^2+^ spike in *pmi4* mutant lines was not caused by variation of aequorin expression across different transgenic lines.

**Figure 5.**
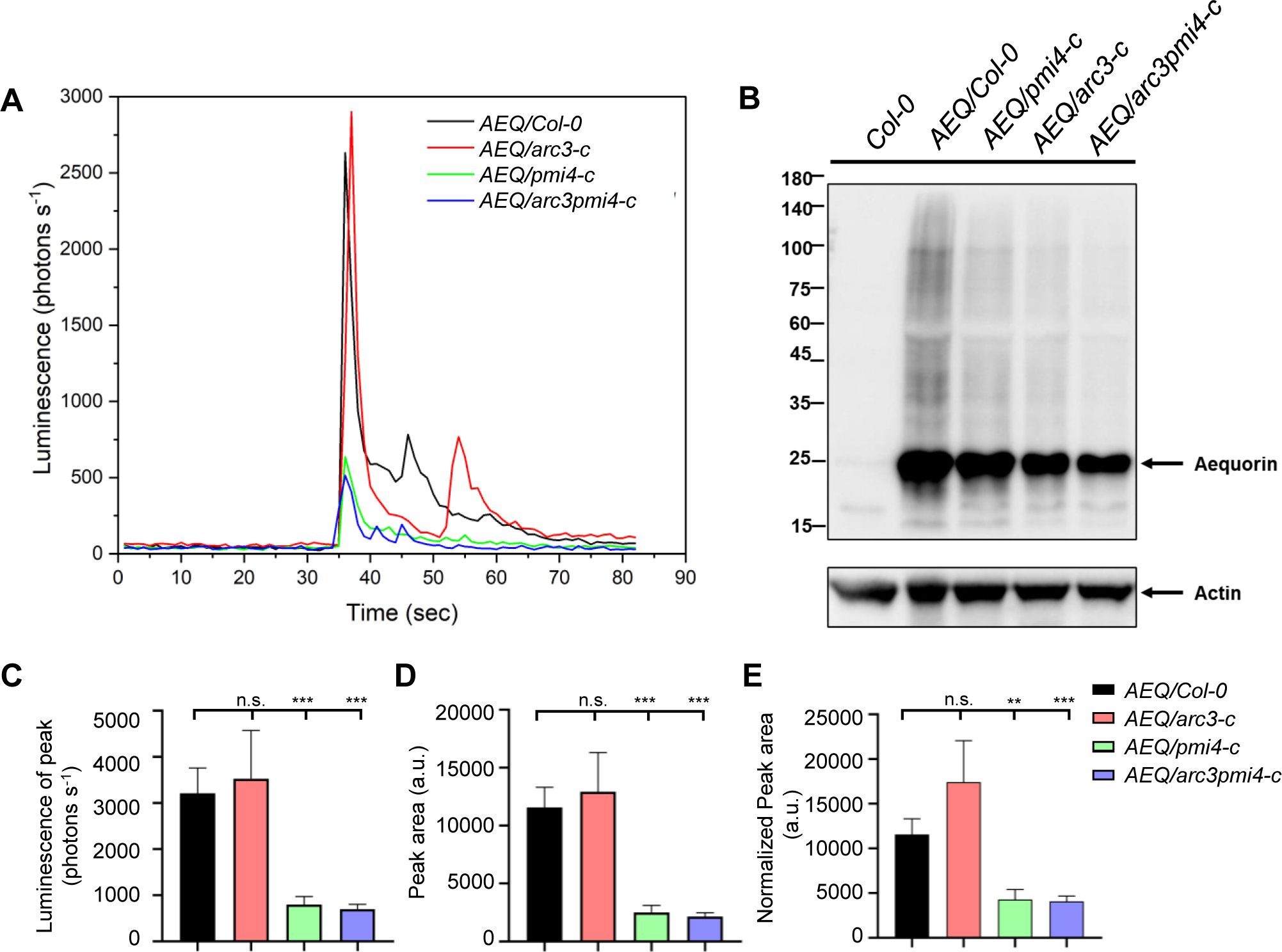
*AEQ/pmi4-c* mutant exhibited an impaired touch-induced rapid Ca^2+^ influx. (A) Representative touch-induced Ca^2+^ influx pattern in *AEQ* reporter line (*AEQ/Col-0*) and corresponding CRISPR-generated *AEQ/arc3-c*, *AEQ/pmi4-c* (*AEQ/pmi4-c1*), *AEQ/arc3pmi4-c* mutant lines. (B) Aequorin expression level was checked using immunoblot. Anti-aequorin antibody was used. (C) Comparison of peak value among the tested mutants. (D) Comparison of peak area among the tested plants. (E) Comparison of peak area normalized with aequorin expression level among the tested plants. Significance among tested plants (n > 30 for each genotype) for touch response was obtained using student t-test: (***, P values less than 0.001; **, P values less than 0.01; *, P values less than 0.05; n.s., not significant).

To further explore the role of PMI4 in touch signaling over the long term, we also investigated the impact of the *pmi4* mutant on touch-induced transcriptomic alterations. The cellular plant force response also can be classified into <u>P</u>rompt <u>F</u>orce <u>R</u>esponse (PFR, the response takes place between 1 and 5 minutes (14)) and <u>R</u>apid <u>F</u>orce <u>R</u>esponse (RFR, the response happens between 5 minutes and 3.5 hours). Thus, in this transcriptomics study, we performed RNA-seq on both *Col-0* and *pmi4* mutant for two different periods of automated hair touch treatments, *i.e* plants in one group were touched for 3 minutes, which belongs to plant prompt force response (PFR), and plants in the other group were touched for 3 minutes and then waited for 17 minutes before collecting with liquid nitrogen, which belongs to rapid force response (RFR) (see Material and Methods part, “Classification of different force-responsive phases”). As a result, 195 and 148 *pmi4*-suppressed (Log_2_ (*Col-0/ pmi4*) > 0.4, q-value <0.05) and *pmi4*-suppressed (Log_2_ *(Col-0/ pmi4*) > 0.4, q-value <0.05) DEGs (Differentially Expressed Genes) were identified for plant PFR and RFR, respectively (Supplemental Table S3a, S3b, Supplemental Fig. S13A, S13B). In addition, 38 and 53 DEGs of these DEGs were previously identified already as “Core touch-responsive genes (13)” in PFR and RFR DEGs, with more than 90% of those DEGs distributed in the *pmi4*-suppressed DEG group (Supplemental Table S3a, S3b), suggesting that *pmi4* suppressed the touch responsive gene expression in both PFR and RFR phases. Comparing with the composition of two groups of DEGs identified from both PFR and RFR, the ratio of *pmi4* -suppressed/ *pmi4* -enhanced showed no significant difference, whereas the “Core touch-responsive genes” is much more abundant (1.83-fold) in the RFR phase (Supplemental Fig. S13C, S13D).

We further analyzed the PFR to RFR transitioning (RFR/PFR) DEGs in *Col-0* and *pmi4*. As a result, 1440 and 1573 RFR/PFR DEGs (q-value <0.05) were identified in *Col-0* and *pmi4*, respectively, with 1029 common RFR/PFR DEGs (Supplemental Table S3c). Subsequently, we compared the 1029 common DEGs with altered touch-responsive expression in both *Col-0* and *pmi4*, with 816 of those DEGs previously identified as “Core touch-responsive genes” (Supplemental Table S3c). Ratiometric comparison selected 58 *pmi4 -*suppressed RFR/PFR DEGs with ΔLog_2_Ratio (*Col-0* - *pmi4*) ≥ 0.4, and 30 *pmi4* enhanced RFR/PFR DEGs with ΔLog_2_Ratio (*Col-0* - *pmi4*) ≤ -0.4 were selected (Fig. 6A). We then focused on the *pmi4* -suppressed RFR/PFR DEGs with larger difference (ΔLog_2_Ratio (*Col-0* - *pmi4*) ≥ 0.6) with GO enrichment analysis (Fig. 6B, Supplemental Table S3c). The result suggested that those transcripts are related to several phytohormones JA, ET, and ABA, as well as cellular response to oxygen level (Fig. 6B). Comparing the *pmi4* -suppressed (Log_2_ (*Col-0/ pmi4*) ≥ 0.4 in PFR and RFR, and ΔLog_2_Ratio (*Col-0* - *pmi4*) ≥ 0.4 in RFR/PFR ratiometric comparison) “Core touch-responsive genes”, 6 genes were found in the RFR phase and RFR/PFR intersection, including well-known touch responsive gene TCH4 (Supplemental Fig. S13E).

**Figure 6.**
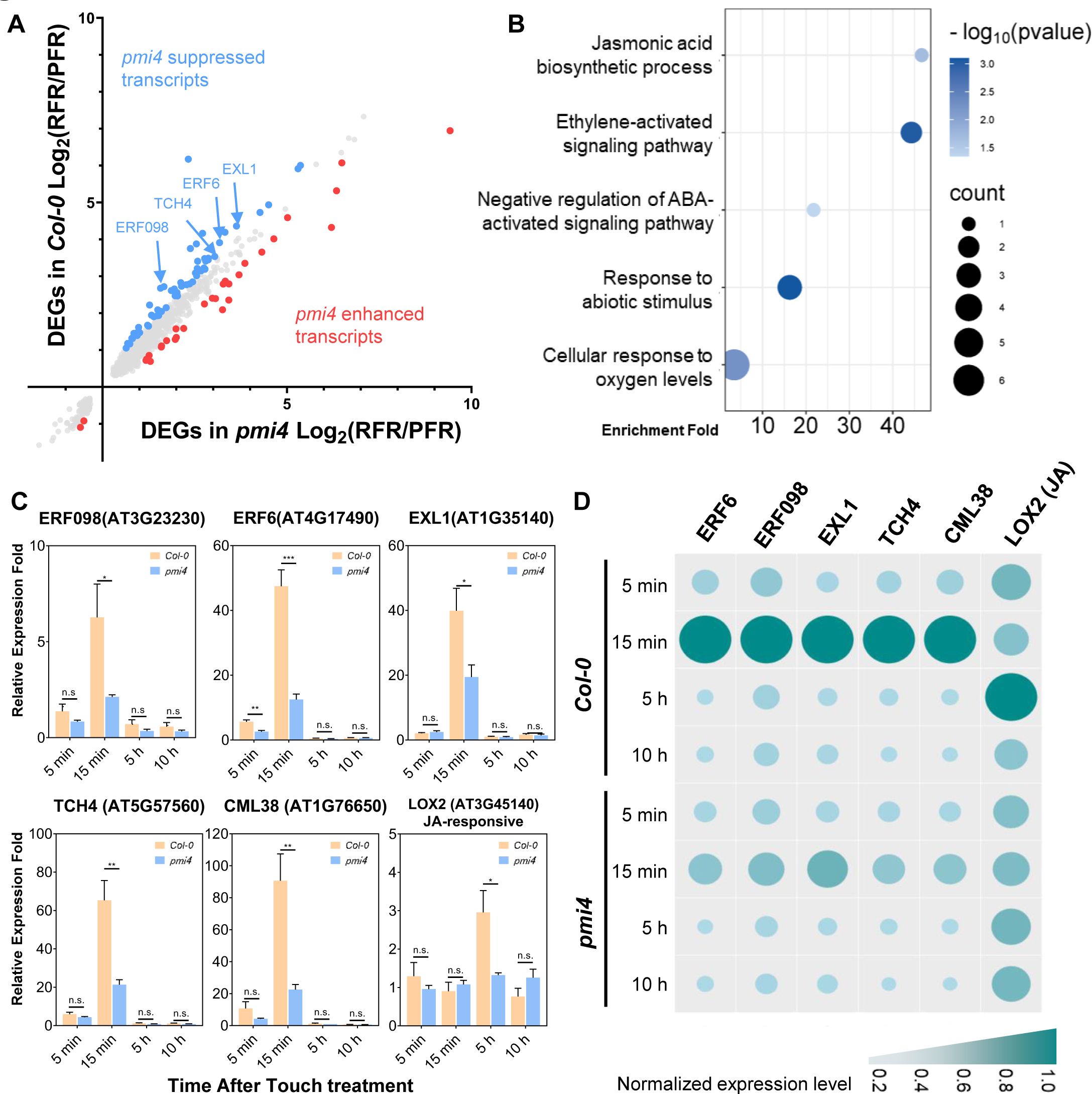
Transcriptome analysis of *pmi4* mutant and wild-type *Col-0* plants at different force-responsive phases. (A) Ratiometric comparison of RFR-PFR DEGs between *pmi4* (x-axis) and wild-type *Col-0* (y-axis). Blue dots indicate 58 *pmi4* suppressed RFR/PFR DEGs with Δlog_2_Ratio (wt - *pmi4*) ≥ 0.4, red dots indicate 30 *pmi4* enhanced RFR/PFR DEGs with Δlog_2_Ratio (wt - *pmi4*) ≤ 0.4. The arrow indicates several DEGs selected for further RT-qPCR analysis in this study. Data can be found in Supplemental Table S3c. (B) GO analysis on the biological process of 23 transcripts with Δlog_2_Ratio (wt - *pmi4*) ≥ 0.6, with the x-axis indicating enrichment fold, y-axis indicating different terms, node size indicating number of protein enriched, and color gradient indicating p-value from Fisher’s exact test. (C) Relative expression level of *Col-0* and *pmi4* at different time points after touch treatment on four DEGs (ERF098, ERF6, EXL1, TCH4), one published touch-induced gene (CML38), JA-responsive transcripts. The mRNA levels were quantified using RT-qPCR by the 2^-ΔΔC^ method. Mean ±SD of four biological repeats is shown. St nt’ t-test was used to analyze the statistics: *, **, ***, and n.s. indicate p value < 0.05, p value < 0.01, p value < 0.001, and not significant. Original data can be found in Supplemental Table S3d. (D) Bubble heatmap showing the relative expression level of selected DEGs and hormone-responsive genes from RT-qPCR result. The relative expression level was normalized (0 - 1) for each gene. Original data can be found in Supplemental Table S3d. Relating supplemental figures can be found in Supplemental Fig. S14. The T-DNA insertion mutant of *pmi4* was used in this transcriptome analysis.

To further validate the possible signaling components that are supposed to be suppressed by *pmi4* mutation, we did RT-qPCR analysis on 4 selected *pmi4*-suppressed DEGs (*ERF6*, *ERF098*, *EXL1*, and *TCH4*) from ratiometric comparison results of RFR/PFR DEGs (Supplemental Table S3c), 1 DEG from the previous touch-induced transcripts, *CML38* (12), and 4 previously reported hormone-responsive DEGs, *LOX2 (*Lipoxygenase 2) (*51*), *AtCP1 (*Cysteine proteinase 1) (*52*), *RD20 (*Responsive to desiccation 20)(*53*), *AGROS (*Auxin-regulated gene involved in organ size) (*54*). The results showed that *pmi4* suppressed all 5 touch-induced DEGs as compared to that of *Col-0* in rapid force response (RFR) (Fig. 6C, Supplemental Fig. S13F). As for the hormone-responsive genes in the <u>S</u>low <u>F</u>orce <u>R</u>esponse (SFR, the response occurs after the force has been applied for more than 3.5 hours) phase, those selected ET and GA responsive genes *AGROS* and *AtCP1* showed no significant change upon touch treatment in three force response phases (PFR, RFR and SFR) investigated (Supplemental Fig. S13F), while JA and ABA-responsive genes *LOX2* and *RD20* were induced to 2.95-fold and 17.06-fold at 5 hours (SFR) in *Col-0* (Fig. 6C-D, Supplemental Fig. S13F), respectively. RT-qPCR also showed that *pmi4* suppressed the induction of JA-responsive gene *LOX2* at 5 h compared to *Col-0* (Fig. 6D).

## Discussion

The plastid movement has been demonstrated to be a trigger for the internal force signaling during gravity-sensing (55) while in the case of cell signaling mediating the external forces (such as touch), there are three theories describing the possible initial force-sensing mechanism, *i.e.* the stretch-activated channels (6), intracellular turgor pressure and subsequent cytoplasmic streaming (7) and cell wall-plasma membrane-cytoskeleton continuum (7, 56). In this study, we have performed the functional proximity labeling-based biotinylproteomics on WPRa4, a member of plastid movement-related protein family which was characterized as touch-responsive signaling components (12). The discovery of 20 putative interactors as well as the subcellular localization study of WPRa4, led to the study of the role of plastid in *Arabidopsis* mechanoresponse. Finally, of special interest was the finding that the *pmi4* mutant plant defective in plastid stroma-localized and tubulin-like plastid protein strongly suppressed the touch-responsive Ca^2+^ transient and touch-inducible transcripts (Fig. 5 and Fig. 6, Supplemental Fig. S13, Supplemental Table S3).

As multiple mechanosensitive channels were identified in Arabidopsis, exploration of ion channel mutants in touch-induced Ca^2+^ influx and thigmomorphogenesis response has been reported in the past few years. Given that the well-known mammalian Pizeo ion channel homolog PIEZO1 in Arabidopsis was recently found to be involved in root mechano-sensing (57–60), we indeed compared the bioluminescence signal of Ca^2+^ transient emitted from the transgenic plant *AEQ/pizeo1* (Supplemental Fig. S12A), we found that *AEQ/pmi4-c* suppressed calcium transient while the *AEQ/piezo1* mutant did not inhibit on the touch-induced calcium transient (Supplemental Fig. S12B). On the other hand, we speculated that the touch-induced Ca^2+^ signal may originate from the plastid and function as a retrograde signal. It has been reported that the plastid Ca^2+^ reservoir is also known to play a role in MAPK signaling activation through the plastid calcium sensor protein CAS (61). There are multiple mechanosensitive ion channels reported in Arabidopsis in the past few years, supporting the stretch-activated mechanosensitive channels’ role as plant mechanosensory (6). Our data support the hypothesis with the plastid as part of the Ca^2+^ supplier. WPRa4-TID based PL-proteomics identified the plastid translocon TIC-TOC complex subunit Tic100 as a putative interactor. It is known that the TIC-TOC complex spans across both inner and outer membranes, the loss-of-function mutant of certain subunit causes defective plastid division and protein import (62). Furthermore, Keissling et al proposed the ‘plastoskeleton’ after observing the plastid-specific filamentous scaffolds with FtsZ network in moss plastid (63) and they further reported several complex morphologies (64). Taken together, based on these reports and our new findings, we believe that the PMI4-comprised plastoskeleton connected with part of cytoskeleton consisting of several WPR family proteins may be organized into the unique touch-responsive structural biomolecular continuum, functioning upstreaming of both the Ca^2+^ transient, which probably originates from plastids, and protein phosphorylation triggered from WPR proteins-linked and membrane-localized receptor-like kinases (such as WR-RLK1, Supplemental Fig. S14, Supplemental Table S2b) to elicit the mechano-signaling in Arabidopsis plant.

Cellular force response to mechano-stimulation is developed in a time-dependent manner (65). In this study, we further characterized this process into four different phases based on the order of occurrence outlined in previous studies (see Materials and Methods for details) and summarized our results in the new model (Supplemental Fig. S14). Touch-induced transcriptome alteration has been reported to occur as early as 5 minutes (14), while more than 92 % percent of DEGs returned to untreated transcript levels after 3 hours (13). With the comparative transcriptome analysis and RT-qPCR validation, multiple touch-responsive transcripts were found to be suppressed in *pmi4* mutants at both PFR and RFR phases. Nonetheless, we evaluated the possible effect on phytohormone regulation in *pmi4* mutants using selected phytohormone-responsive genes. The JA synthesis-related lipoxygenase LOX2 induction was found to be blocked by the *pmi4* mutant during the SFR phase, indicating that *pmi4* insensitivity toward touch force is related to the JA regulation during thigmomorphogenesis. The role of JA in promoting thigmomorphogenesis was well characterized by (15) using the JA-deficient mutant *aos* and JA-constitutive expression transgenic plant *OPR3-OE*. As JA biosynthesis precursor 12-oxophytodienoic acid (OPDA) relies on plastid lipid as substrate and LOX2 as crucial lipoxygenase for this process (66), the effect of *pmi4* on the JA signaling pathway served as a much more reasonable explanation of its touch insensitivity. A recent study highlighted the crosstalk of multiple phytohormones under mechanical stress (14), further exploration of the role of plastid in phytohormone hemostasis could provide unique clues on the organelle-mediated abiotic stress response.

Based on data presented in this study and previous studies, a new model is therefore proposed here, in which the cytoskeleton-plastoskeleton continuum linked with unknown plastidic calcium channels might serve as the initial force-sensing network (Supplemental Fig. S14). Upon touch stimulation, both the plastoskeleton protein PMI4 and cytoskeleton protein PMI proteins, including WPRa4 protein, receive the touch force signal generated from the cytoplasmic pressure and the plastid movement alteration, which in turn triggers the Ca^2+^ transient, or presumably the retrograde calcium oscillation from plastids, to activate downstream phosphorylation of CPKs and CAMTA3 (12, 22). Consequently, the phosphor-relay on these signaling proteins transduces the force signals into the nucleus to activate the touch-responsive genes (Supplemental Fig. S14). Taken together, the multidisciplinary work including the PL-based PTM proteomics on WPRa4, the thigmomorphogenesis study as well as the Ca^2+^ transient and transcriptome analysis on *pmi4* mutants strongly suggested a role for the cell wall-plasma membrane-cytoskeleton-plastid continuum as the early steps of sensing the mechano-stimulation and in initiating the touch-triggered cell signaling (Supplemental Fig. S14). It is therefore postulated that, upon touch stimulation, both the plastidic PMI4 protein networks receive the touch force signal generated by either plastid physical extrusion, which also serves as part of the Ca^2+^ pool for touch-induced Ca^2+^ transient during the IFR phase. A possible linkage might exist between WPRa4 and PMI4 protein skeletal networks either through TIC-TOC complex (PL data from this study) or through PAP85 (a vicilin-like protein involved in pathogen defense response (67), XL-MS data from our published dataset (36)). Phosphorylation on several signaling components, including MKK1/2, WPRa4, and CAMTA3 (12, 22) subsequently initiated the signal transduction starting from the IFR phase and prolonged to the PFR phase of touch response. Transcription of touch-inducible genes starts from the PFR phase and matures during the SFR phase, lasting for several hours and affecting further morphological changes. The whole model describes a systematic molecular network organized for plant force response.

## Supporting information

Manuscript

Manuscript

Manuscript

Manuscript

## ABBREVIATIONS

XIC: Extracted Ion Chromatogram
MS: mass spectrometry
SQUA: stable isotope-based quantitation
Col-0: Columbia-0
TID: TurboID
TREPH1: Touch-Regulated Phosphoprotein 1
WEB1: weak chloroplast movement under blue light 1
PMI1: plastid movement impaired 1
PMI2: plastid movement impaired 2
WPRa4: WEB1/PMI2 Related protein family member a4
WR-RLK1: WPRa4-related receptor-like kinase 1
Tic100: Translocon at the inner envelope membrane of chloroplasts 100
FtsZ1: Arabidopsis Thaliana homolog of bacterial cytokinesis z-ring protein
WIRK1: WInd-Related Kinase 1
TCH4: TOUCH 4
GO: Gene Ontology
CML38: Calmodulin-like 38
EXL1: Exordium like 1
ERF6: Ethylene responsive element binding factor 6
ERF98: Ethylene responsive element binding factor 98
LOX2: Lipoxygenase 2
AtCP1: Cysteine proteinase 1
RD20: Responsive to desiccation 20
AGROS: Auxin-regulated gene involved in organ size
UEB: urea protein extraction buffer
F: forward mixing of peptide samples
R: reciprocal mixing of peptide samples
DDA: data-dependent acquisition
PSM: peptide spectrum match
HCD: higher-energy collisional dissociation
FDR: false discovery rate
SD: Standard deviation

## Acknowledgments

These authors want to send special thanks to Prof. Alice TING of Stanford University for sharing the original TurboID plasmid, Dr. Marc Knight for the AEQ gene, Dr. Thibaud for the plasmid of G5A and G5A/Col-0 transgenic plant, Prof Ardem Patapoutian for the piezo seeds in pNano system, and Prof. Osteryoung for sharing the T-DNA mutant of *pmi4, arc3,* and *arc3pmi4*. Special thanks to Dr. Autar Mattoo for his generous and insightful advice on the manuscript preparation. We also express our gratitude to Dr. Daiying XU and Jingjing Li of NanoBioImaging Ltd., Dr. Yan Zhang of Materials Characterization and Preparation Facility, HKUST. These authors want to express their great thanks to Dr. Junxian He and his PhD student Mr. Wei Yang for their kind help with the confocal microscopy imaging. We also thank Dr. Clarence CHENG and the Biosciences Central Research Facility (BioCRF) proteomic core of HKUST for their support in handling the MS sample analysis.

## Data Availability

All study data are included in the article and/or supporting information. The mass spectrometry proteomics data have been deposited to the ProteomeXchange Consortium via the PRIDE partner repository with the dataset identifier of PXD055777. Reviewers can access these original datasets with the following account (Username, Password: jg2HKXc2nTGW).

## Author Contributions

K.W. and M.L contributed to the *WPRa4-TID* and *TurboID* transgenic plants-making; K.B.W. performed the quantitative biotinylprotemics on *WPRa4-TID* and *TurboID* transgenic plants and bioinformatic analysis of proximity-labeling PTM proteomics results; K.W. and S.L. performed the microscopic analysis; N.Y. conducted the Ca^2+^ oscillation experiments and transcriptomics. N.Y. and K.B.W. conducted transcript quantitation using RT-qPCR for both touched *pmi4* and *Col-0*; K.B. W. performed the plastid isolation and immunoblot analysis. N.Y. and J.R performed the touch response assays; S.D. performed the XL-MS; N.L., K.B.W., N.Y. wrote the manuscript; N.L., Z.Y., Y.L., S.D., Y.A., F.T., Z.G., Y.Z., and W.Y., contributed to the revision of manuscript and the supporting to and discussion on experiments; N.L. was in charge of project planning, designing, project execution, supervision of experiments, and communication with collaborators and responsible for distribution of materials integral to the findings.

