## Supplementary material for "Functional, Biotinylproteomic and Bioinformatic Analysis of Both Cytoskeletal and Plastoskeletal Proteins in Plant Mechanoresponse": Manuscript

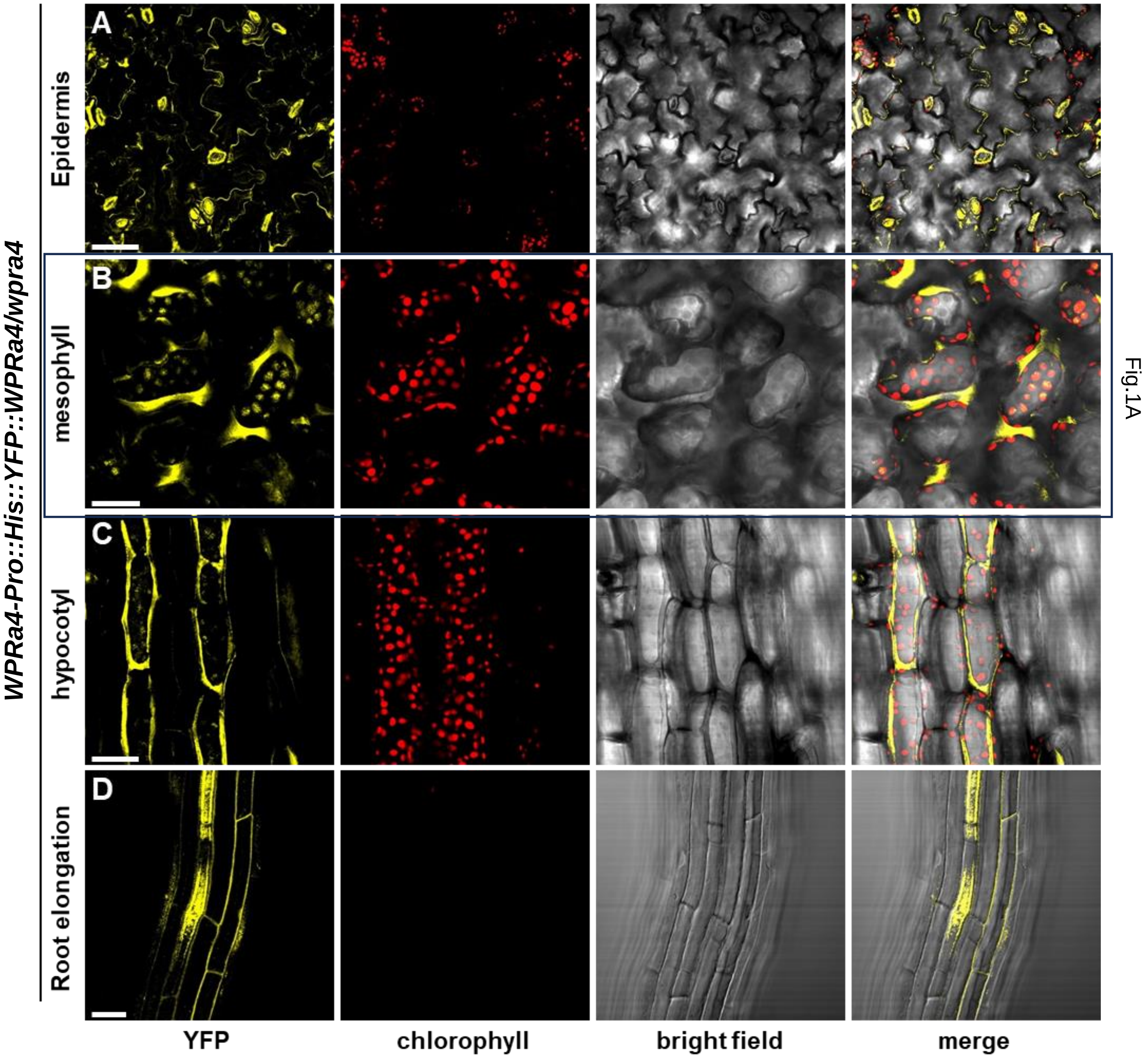


**Figure S1. Confocal Microscopy study of YFP-WPRa4 localization under native promoter.**

(A) to (D) Subcellular distribution of WPRa4 from *Pro::His::YFP::WPRa4/wpra4* transgenic plant in (A) epidermis, (B) mesophyll (also shown in Fig.1a), (C) hypocotyl, and (D) root elongation cells. Scale bars are 60 μm (A) and 30 μm (B to D). The excitation of YFP and chlorophyll auto-fluorescence were performed with 515 nm and 635 nm argon lasers, respectively. The detected emission ranges of YFP and chlorophyll auto-fluorescence were 530 – 570 nm and 655 – 755 nm, respectively.

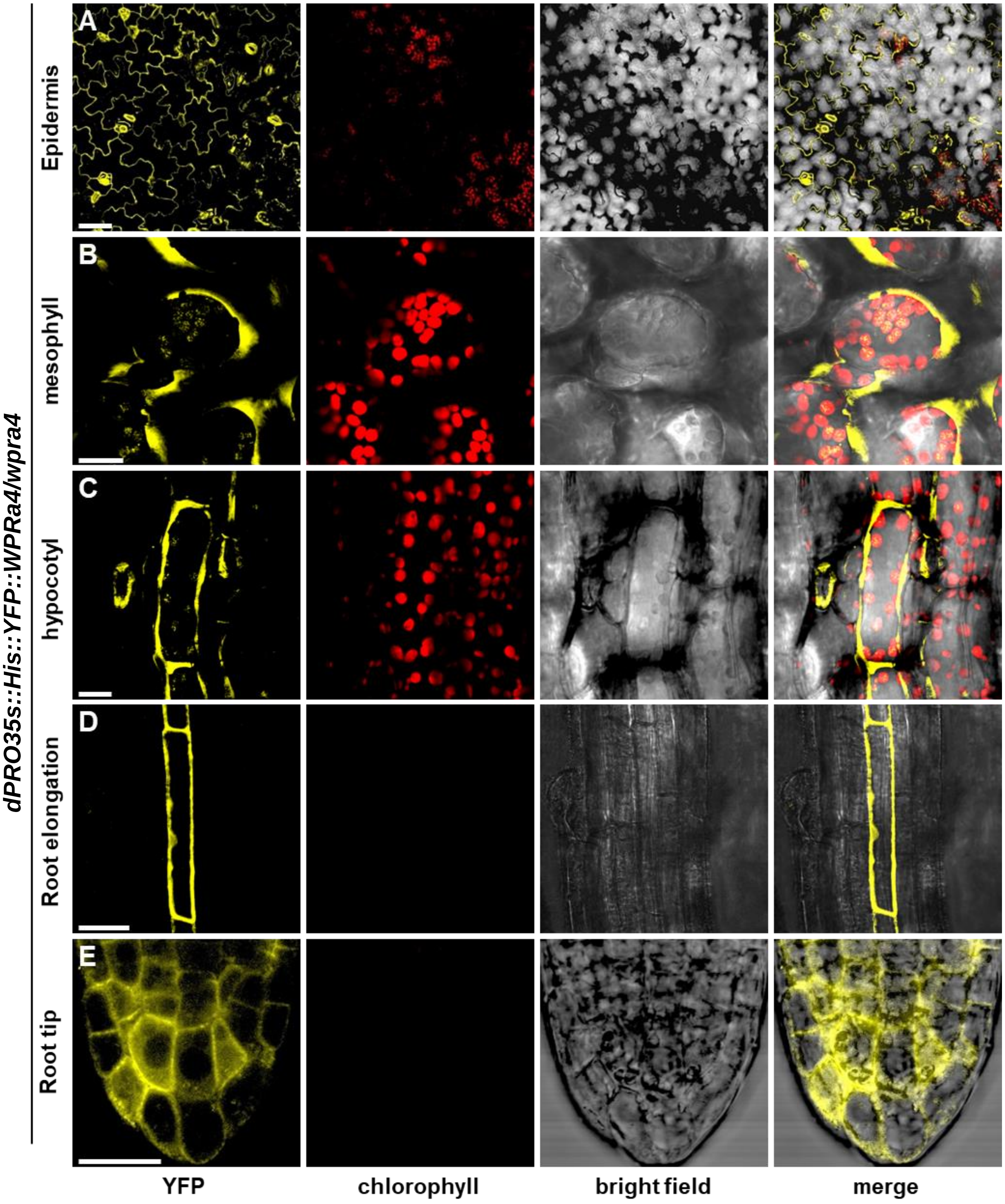


**Figure S2. Confocal Laser Microscopy imaging of the subcellular distribution of WPRa4 fusion protein.** (A) to (E) Subcellular distribution from *dPro35S::His::YFP::WPRa4/wpra4* transgenic plant in epidermal (A), mesophyll (B), hypocotyl (C), root elongation (D) and root tip (E) cells, respectively. Scale bars are 60 µm (A), 30 µm (B to D), and 20 µm (E). The excitation of YFP and chlorophyll auto-fluorescence were performed with 515 nm and 635 nm argon lasers, respectively. The emission ranges of YFP and chlorophyll auto-fluorescence were 530 – 570 nm and 655 – 755 nm, respectively. This figure is related to Fig. 1a.

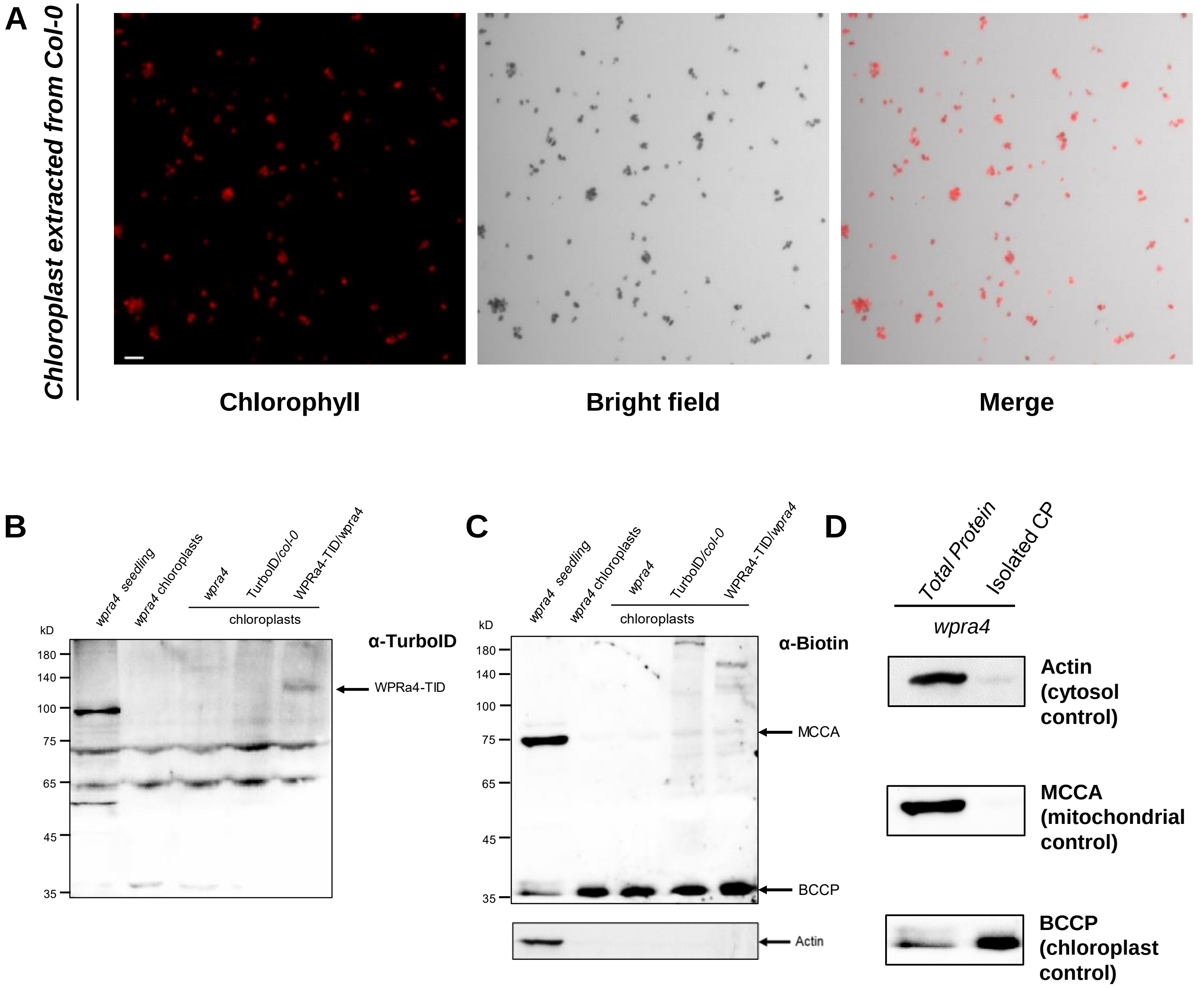


**Figure S3. Extraction of chloroplasts and immunoblot confirmation of the presence of WPRa4-TID on chloroplasts.** (A) Chloroplast extraction purity checking from wild-type *Col-0* plant. (B) TurboID fusion protein expression pattern of chloroplast proteins from TurboID harboring transgenic plants. Anti-TurboID antibody was used for the immunoblot. The letter α stands for 'anti'. (C) Biotinylated protein pattern of 17 days seedlings and chloroplast proteins from TurboID harboring transgenic plants. The membrane is stripped after Fig.S3b and then incubated with the anti-Biotin antibody. Anti-actin for (B) and (C) was applied at the bottom. (D) Cytosol, mitochondrial, and chloroplast control used in this immunoblot. This figure is related to Fig. 1C-1E.

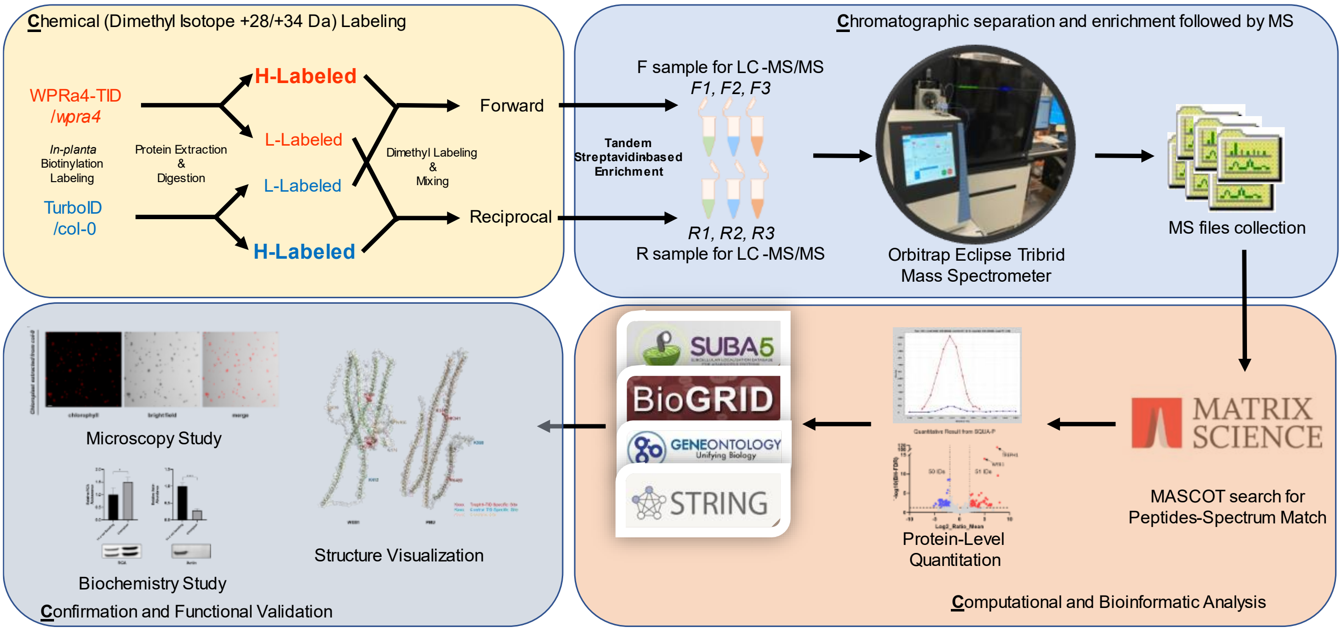


**Figure S4. 4C Biotinylproteomics workflow used in this study.** This method involved four steps: (i) **C**hemical labeling using *in planta* biotin labeling and in vitro peptides dimethyl labeling with $C_{12}H_2$ formaldehyde (light, +28.031 Da) or $C_{13}D_2$ formaldehyde (heavy, +34.063 Da); (ii) **C**hromatographic separation and mass spectrometry/mass spectrometry (MS/MS) identification after high purity enrichment of biotinylated peptides; (iii) **C**omputational and bioinformatics analysis for identification and quantification of WPRa4 proximal proteins and (iv) **C**onfirmation with functional analysis. This figure is related to Fig. 2 and Fig. 3.

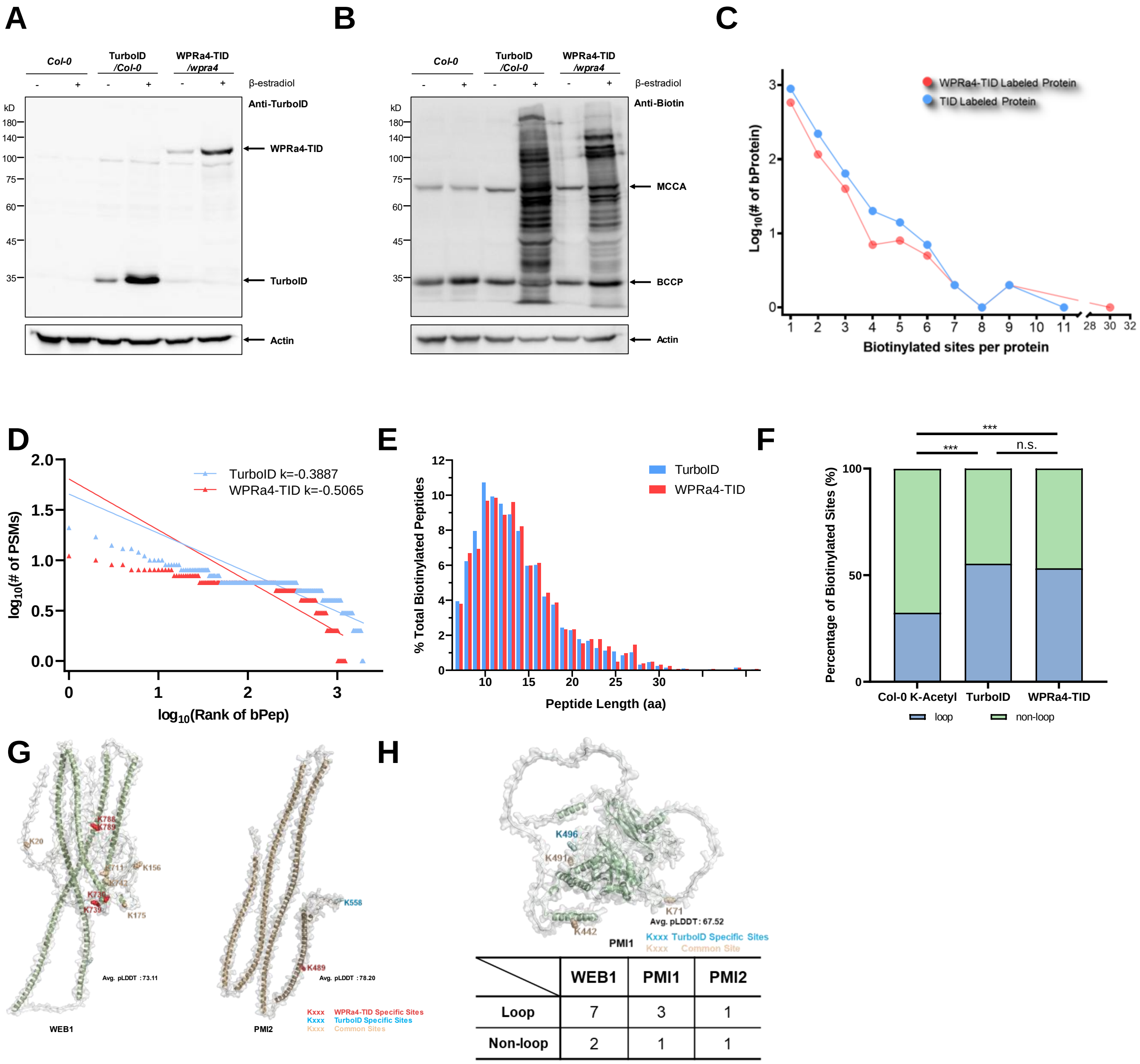


| | WEB1 | PMI1 | PMI2 |
|---|---|---|---|
| Loop | 7 | 3 | 1 |
| Non-loop | 2 | 1 | 1 |

**Figure S5. Supplementary information of WPRa4-TID expression and biotinylated peptides enrichment performance.** (A) β-estradiol induction on *Col-0*, *TurboID*/*Col-0* and *WPRa4-TID*/*wpra4* transgenic plants, using anti-TurboID antibody to detect the expression. (B) β-estradiol induction of biotinylated proteins in *Col-0*, *TurboID*/*Col-0,* and *WPRa4-TID*/*wpra4* transgenic plants, using an anti-biotin antibody to detect the differential biotinylated protein pattern. (C) Summarization of the biotinylated sites on each protein. The Y-axis indicates $Log_{10}$ (Number of proteins), and the X-axis indicates biotinylated sites identified in one protein. TurboID-labeled proteins (Blue), and WPRa4-TID-labeled proteins (Red) were counted. Original data can be found in (Supplemental Table S1c). (D) Summarization of the PSM numbers on each biotinylated peptide. The Y-axis indicates $log_{10}$ (Number of PSMs), and the X-axis indicates biotinylated sites ranking. TurboID control labeled peptide (Blue), and WPRa4-TID labeled peptide (Red) were counted. (E) Summarization of the peptides length distribution. TurboID control labeled peptide (Blue), and WPRa4-TID labeled peptide (Red) were calculated. (F) A bar chart represents a comparison of the percentage of biotinylated sites on loop region or non-loop region of *Col-0* endogenous biotinylated sites, TurboID biotinylated sites, and WPRa4-TID biotinylated sites. Fisher`s exact test was used to compare the difference. The n.s. and * represents $p\text{-value} \geq 0.05$ and $p\text{-value} < 0.05$, respectively (Original data can be found in Supplemental Table S1e). (G) Displaying of biotinylated sites on the AlphaFold predicted WEB1, and PMI2 structures. (H) Displaying of biotinylated sites on the AlphaFold predicted PMI1 structures, with counting of the loop and non-loop sites of WEB1, PMI1, and PMI2. This figure is related to Fig. 2.

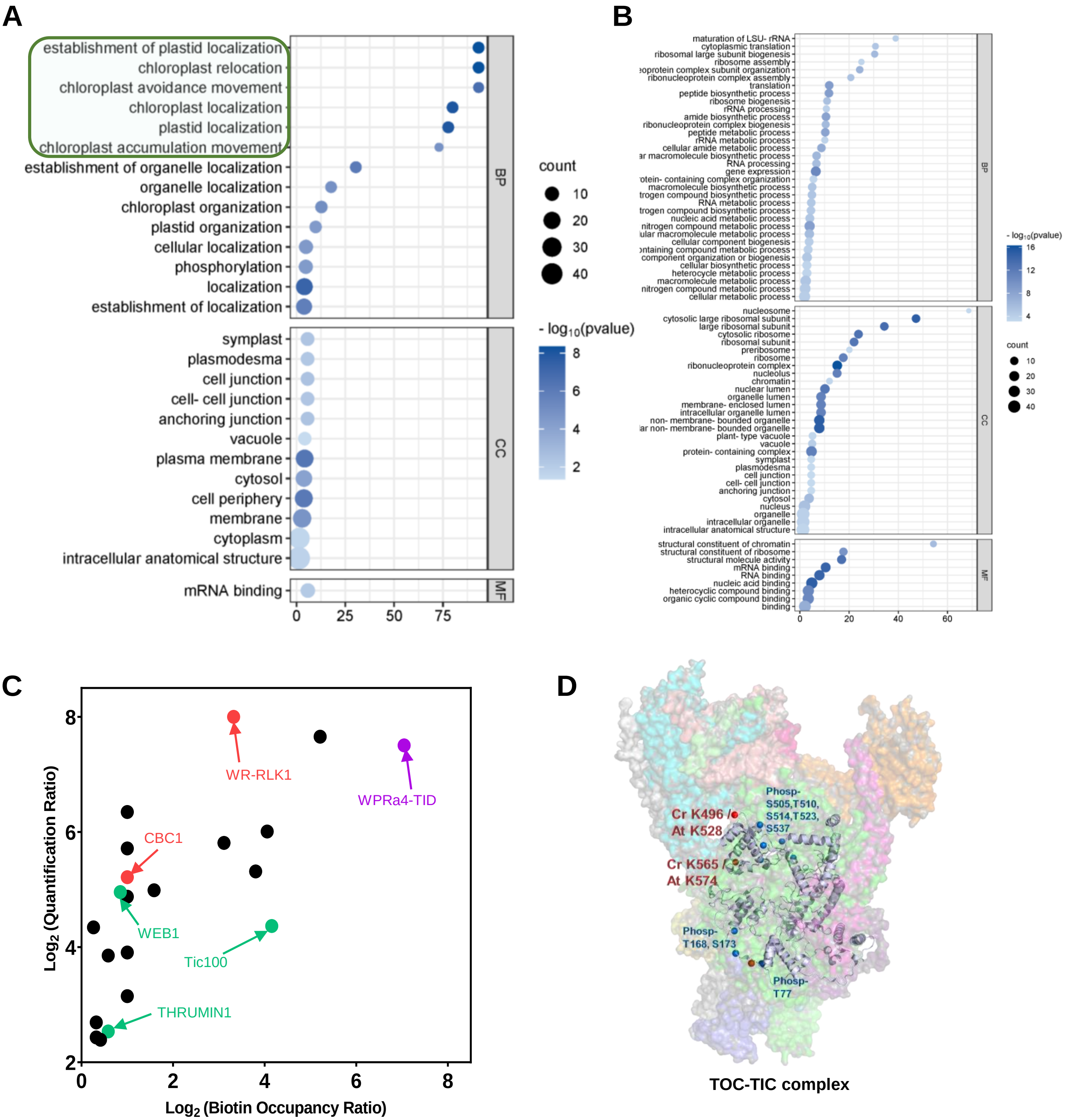


**Figure S6. Bioinformatic analysis related to quantitative biotinylproteomic result.** (A) GO analysis result of the 49 WPRa4 proximal proteins (BP, Biological Process; MF, Molecule Function; CC, cellular component). (B) GO analysis result of the TID proximal proteins (BP, Biological Process; MF, Molecule Function; CC, cellular component). (C) Mapping of 21 significant WPRa4 proximal proteins with quantitative ratio > 2.0, $Log_2(BOR) > 0$. Green dots represent the Plastid-related proteins, red dots represent receptor like kinases, and purple dot represents WPRa4-TID. Data can be found in Supplemental Table S2b. (D) Displaying the WPRa4-specific labeled sites K528 and K574 on Tic 100 from the TIC-TOC complex structure (PDBid: 7VCF). Cr, Chlamydomonas reinhardtii; At, *Arabidopsis thaliana.* This figure is related to Fig. 2 and Fig. 3.

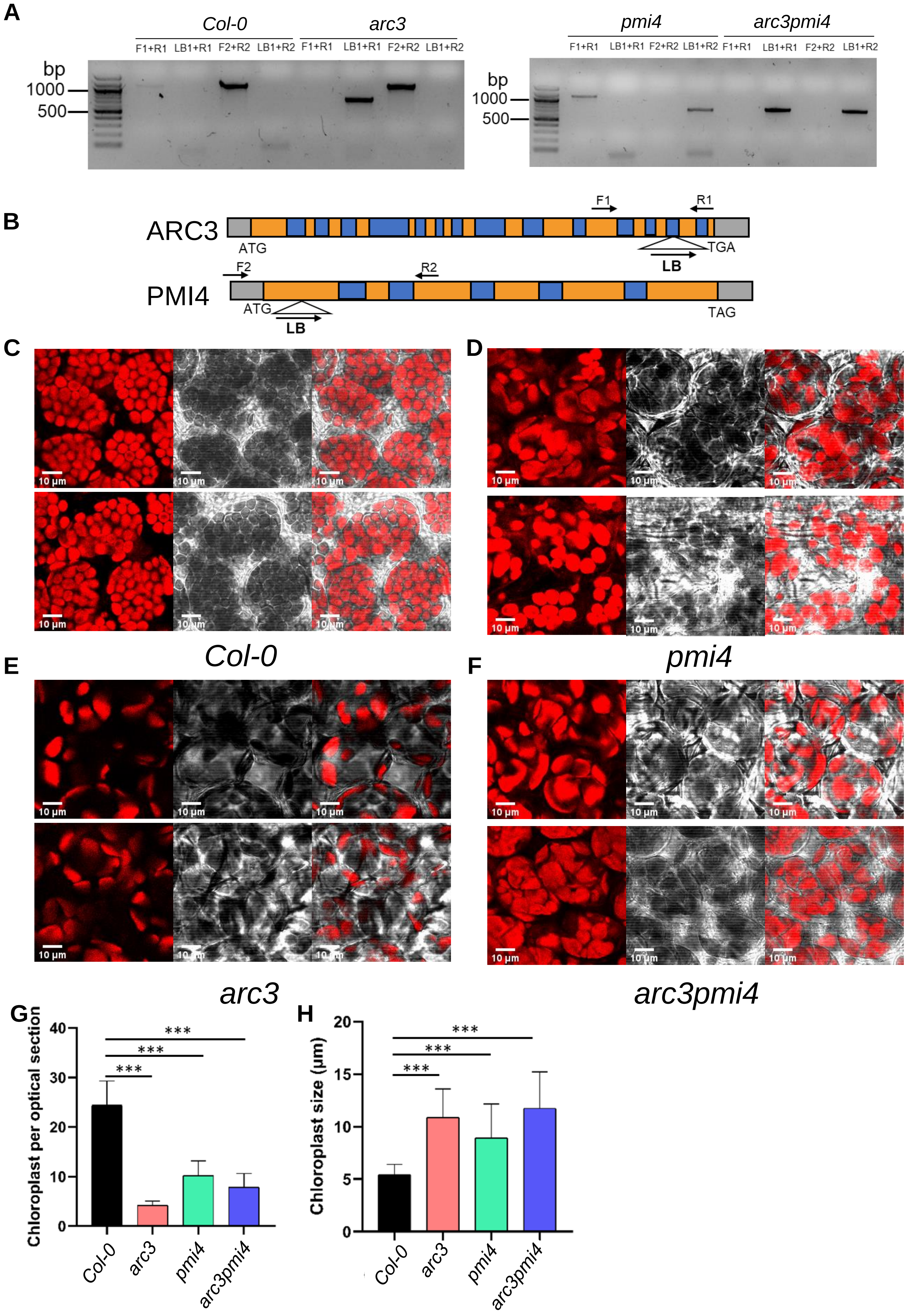
A
Col-0
arc3
pmi4
arc3pmi4
F1+R1
LB1+R1
F2+R2
LB1+R2
bp
1000
500
B
ARC3
PMI4
ATG
TGA
TAG
F1
R1
F2
R2
LB
C
D
E
F
10 µm
Col-0
pmi4
arc3
arc3pmi4
G
Chloroplast per optical section
0
10
20
30
40
***
H
Chloroplast size (µm)
0
5
10
15
20

**Figure S7. Homozygosity and chloroplast morphology examination of *pmi4, arc3, and arc3pmi4* double mutant in this study.**

(A) Gene structure of *ARC3* (AT1G76990) and *PMI4* (AT5G55280). Triangles represent T-DNA insertion sites in T-DNA insertional mutants. Primers used for genotyping are presented with arrows. Grey box, 5`- and 3`-UTR. Orange box, exon region. Blue box, intron region. ATG and TAG/TGA indicate the start and stop codon, respectively. (B) Genotyping of *arc3* and *pim4* mutant using tri-primer method. (c-f) Confocal imaging of chloroplast morphology of (D) wild-type *Col-0*, (D) *pmi4,* (E) *arc3,* and (F) *arc3pmi4*. From left to right: Chlorophyll, Bright Field, Merged image. The scale bar is 10 µm for each image. (G-H) Bar chart showing the chloroplast number per cell (G) and chloroplast size (H) in the above images. Statistical test was performed employing student's t-test with $p < 0.001$ for pairwise t-test shown as ***.

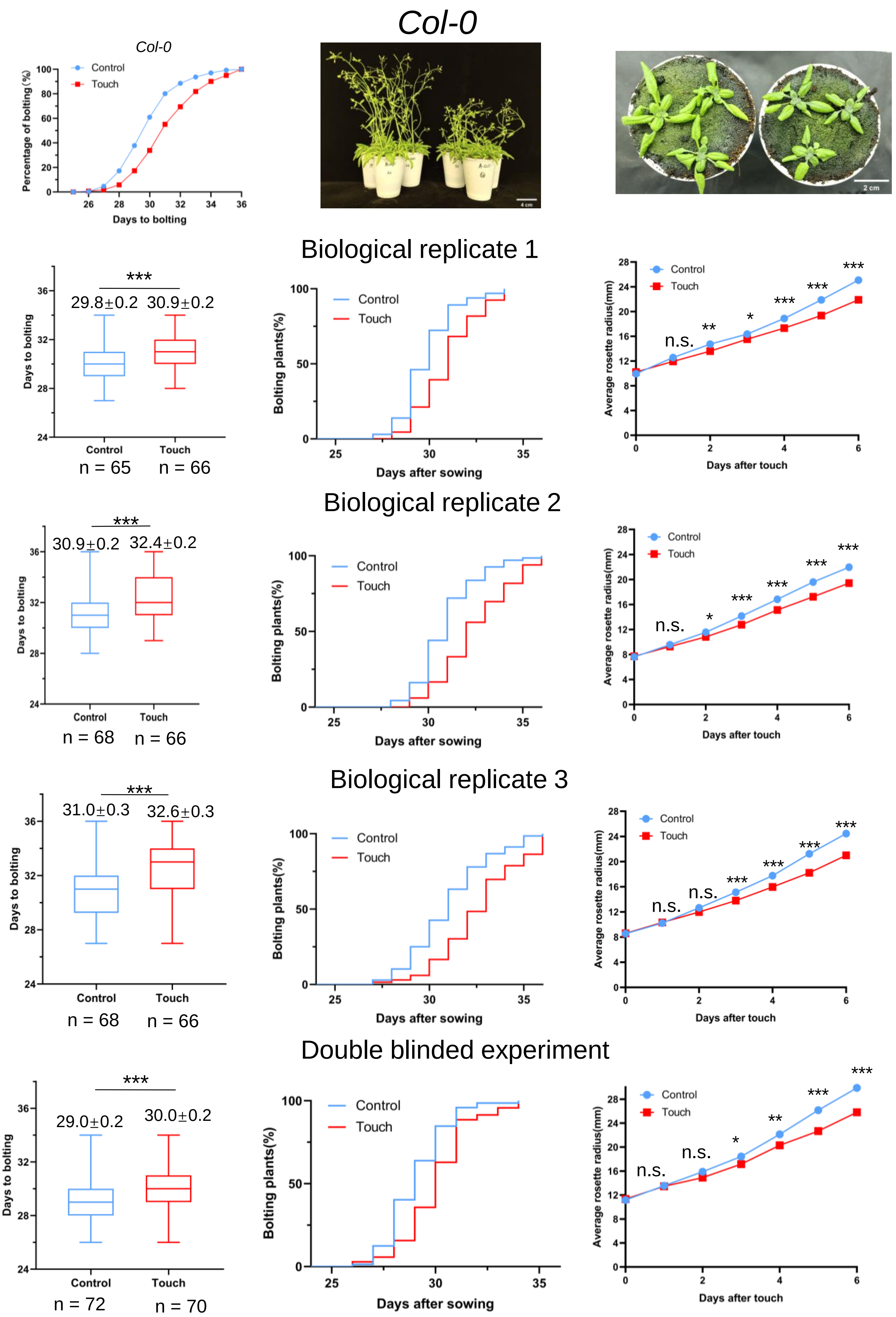
Col-0
Col-0
Control
Touch
Percentage of bolting (%)
Days to bolting
4 cm
2 cm
Biological replicate 1
***
29.8±0.2
30.9±0.2
Days to bolting
Control
Touch
n = 65
n = 66
Bolting plants(%)
Days after sowing
Average rosette radius(mm)
Days after touch
n.s.
**
*
***
Biological replicate 2
30.9±0.2
32.4±0.2
n = 68
n = 66
Biological replicate 3
31.0±0.3
32.6±0.3
n = 68
n = 66
Double blinded experiment
29.0±0.2
30.0±0.2
n = 72
n = 70

**Figure S8. Touch responses of soil-grown wild type (*Col-0*) Arabidopsis to automated human hair touch.** The comparison of bolting plants grown in soil and touched by automated human hair. The above-left panel is the average bolting percentage from all biological experiments. The box and whisker plots of bolting days were analyzed for each biological experiment. The numbers of the total individual plants (n) and means ± SE from 3 biological replicates and one double-blinded experiment are annotated under and above the plots, respectively. The two right-above panels are photographs of a representative of untouched (control) and touch-treated plants (touch) for bolting and plant size. The number of bolting plants was the average bolting number calculated by three people.
Kaplan-Meier plots are shown in the middle, which are the percentage of bolting plants over the growth period (days after sowing). Trend chart plots which are the average rosette radius over the touch period (touch day) are shown in the right panels. Statistical test was performed employing student's t-test. The significance of $p < 0.05$, $p < 0.01$, and $p < 0.001$ for pairwise t-test are shown as *, **, and ***, respectively. This figure is related to Fig. 4.

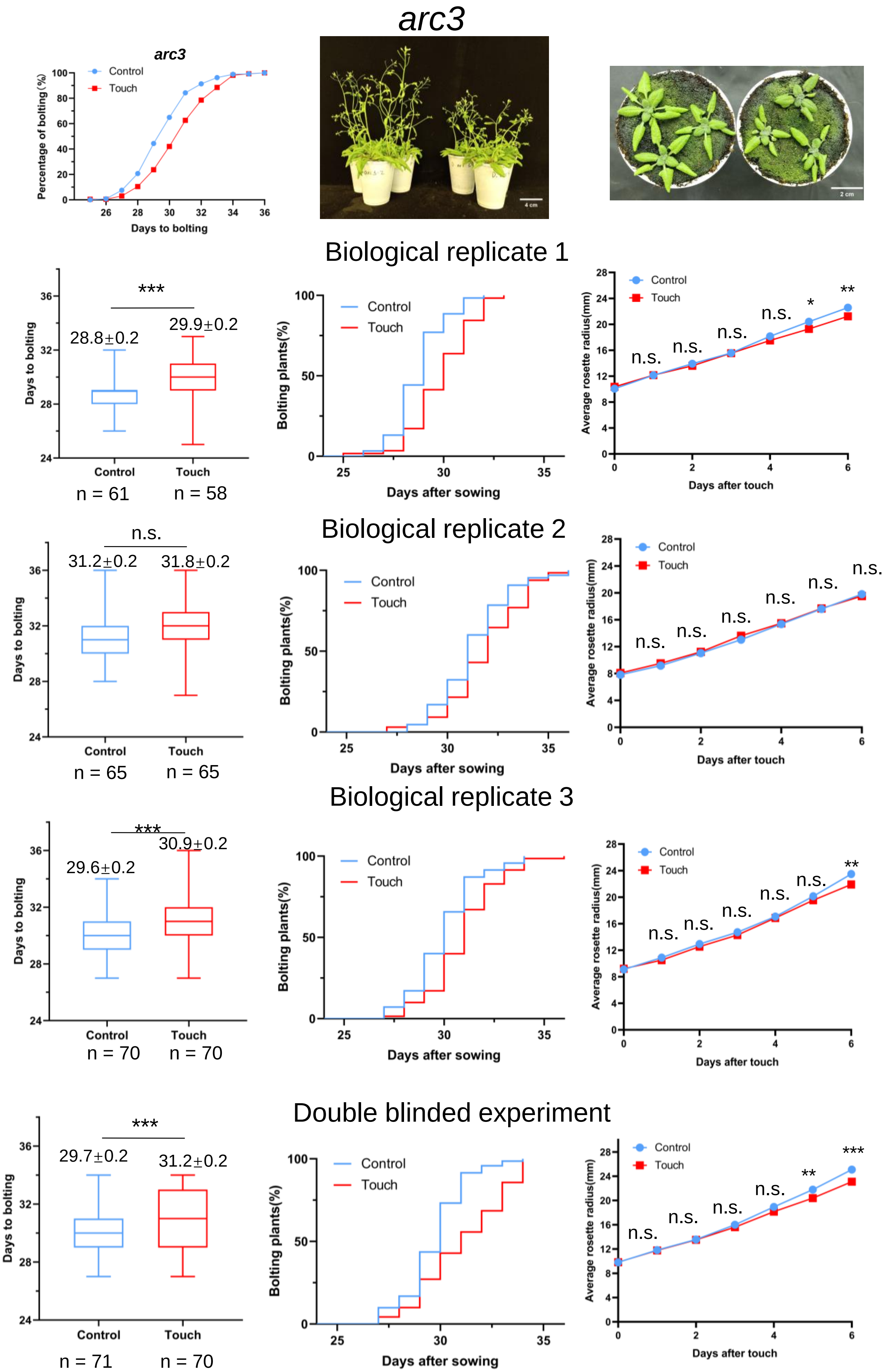
arc3
arc3
Control
Touch
Percentage of bolting (%)
Days to bolting
4 cm
2 cm
Biological replicate 1
***
28.8±0.2
29.9±0.2
Days to bolting
Control
Touch
n = 61
n = 58
Bolting plants(%)
Days after sowing
Average rosette radius(mm)
Days after touch
n.s.
*
**
Biological replicate 2
n.s.
31.2±0.2
31.8±0.2
n = 65
n = 65
Biological replicate 3
***
29.6±0.2
30.9±0.2
n = 70
n = 70
Double blinded experiment
***
29.7±0.2
31.2±0.2
n = 71
n = 70

**Figure S9. Touch responses of soil-grown *arc3* Arabidopsis to automated human hair touch.** The comparison of bolting plants grown in soil and touched by automated human hair. The above-left panel is the average bolting percentage from all biological experiments. The box and whisker plots of bolting days were analyzed for each biological experiment. The numbers of the total individual plants (n) and means ± SE from 3 biological replicates and one double-blinded experiment are annotated under and above the plots, respectively. The two right-above panels are photographs of a representative of untouched (control) and touch-treated plants (touch) for bolting and plant size. The number of bolting plants was the average bolting number calculated by three people.
Kaplan-Meier plots are shown in the middle, which are the percentage of bolting plants over the growth period (days after sowing). Trend chart plots which are the average rosette radius over the touch period (touch day) are shown in the right panels. Statistical test was performed employing student's t-test. The significance of $p < 0.05$, $p < 0.01$, and $p < 0.001$ for pairwise t-test are shown as *, **, and ***, respectively. T-DNA insertion mutant *arc3* was applied in this figure. This figure is related to Fig. 4.

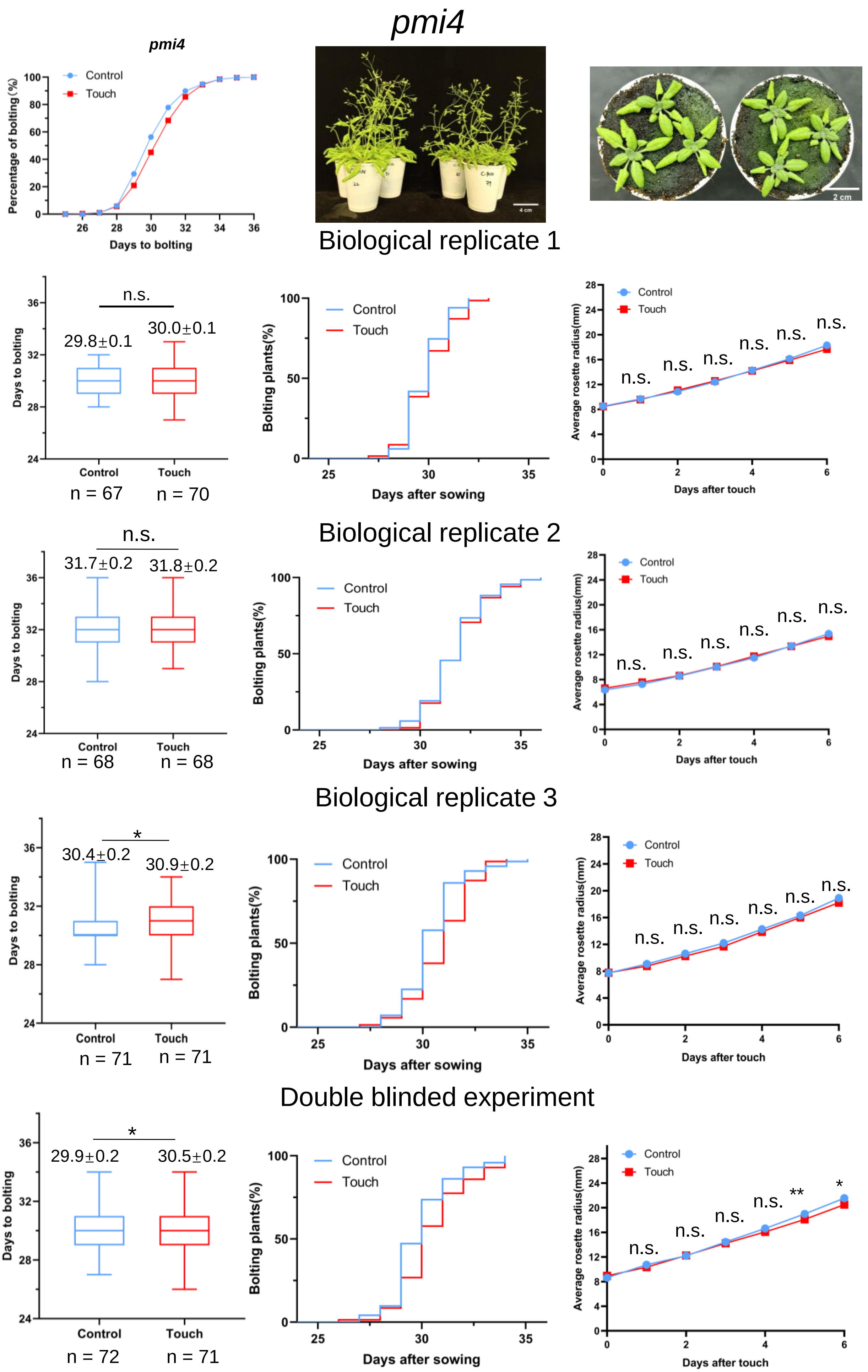
pmi4
pmi4
Control
Touch
Percentage of bolting (%)
Days to bolting
4 cm
2 cm
Biological replicate 1
n.s.
29.8±0.1
30.0±0.1
Days to bolting
Control
Touch
n = 67
n = 70
Control
Touch
Bolting plants(%)
Days after sowing
Control
Touch
Average rosette radius(mm)
Days after touch
n.s.
n.s.
n.s.
n.s.
n.s.
n.s.
Biological replicate 2
n.s.
31.7±0.2
31.8±0.2
Days to bolting
Control
Touch
n = 68
n = 68
Control
Touch
Bolting plants(%)
Days after sowing
Control
Touch
Average rosette radius(mm)
Days after touch
n.s.
n.s.
n.s.
n.s.
n.s.
n.s.
Biological replicate 3
*
30.4±0.2
30.9±0.2
Days to bolting
Control
Touch
n = 71
n = 71
Control
Touch
Bolting plants(%)
Days after sowing
Control
Touch
Average rosette radius(mm)
Days after touch
n.s.
n.s.
n.s.
n.s.
n.s.
n.s.
Double blinded experiment
*
29.9±0.2
30.5±0.2
Days to bolting
Control
Touch
n = 72
n = 71
Control
Touch
Bolting plants(%)
Days after sowing
Control
Touch
Average rosette radius(mm)
Days after touch
n.s.
n.s.
n.s.
n.s.
**
*

**Figure S10. Touch responses of soil-grown *pmi4* Arabidopsis to automated human hair touch.** The comparison of bolting plants grown in soil and touched by automated human hair. The above-left panel is the average bolting percentage from all biological experiments. The box and whisker plots of bolting days were analyzed for each biological experiment. The numbers of the total individual plants (n) and means ± SE from 3 biological replicates and one double-blinded experiment are annotated under and above the plots, respectively. The two right-above panels are photographs of a representative of untouched (control) and touch-treated plants (touch) for bolting and plant size. The number of bolting plants was the average bolting number calculated by three people. Kaplan-Meier plots are shown in the middle, which are the percentage of bolting plants over the growth period (days after sowing). Trend chart plots which are the average rosette radius over the touch period (touch day) are shown in the right panels. Statistical test was performed employing student's t-test. The significance of $p < 0.05$, $p < 0.01$, and $p < 0.001$ for pairwise t-test are shown as *, ** and ***, respectively. T-DNA insertion mutant *pmi4* was applied in this figure. This figure is related to Fig. 4.

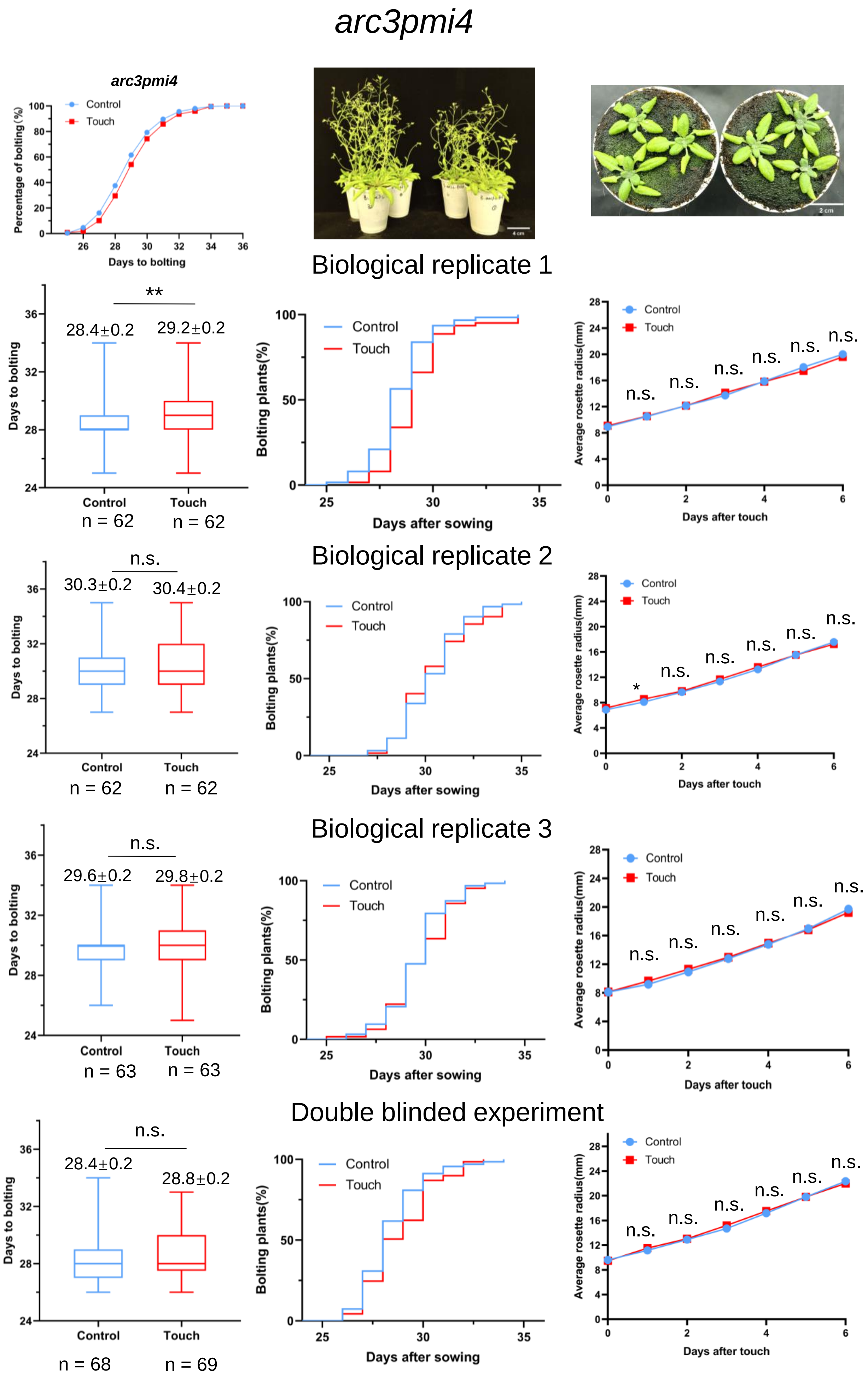
arc3pmi4
arc3pmi4
Control
Touch
Percentage of bolting (%)
Days to bolting
Biological replicate 1
**
28.4±0.2
29.2±0.2
Days to bolting
Control
Touch
n = 62
n = 62
Control
Touch
Bolting plants(%)
Days after sowing
Control
Touch
n.s.
Average rosette radius(mm)
Days after touch
Biological replicate 2
n.s.
30.3±0.2
30.4±0.2
Days to bolting
Control
Touch
n = 62
n = 62
Control
Touch
Bolting plants(%)
Days after sowing
Control
Touch
*
n.s.
Average rosette radius(mm)
Days after touch
Biological replicate 3
n.s.
29.6±0.2
29.8±0.2
Days to bolting
Control
Touch
n = 63
n = 63
Control
Touch
Bolting plants(%)
Days after sowing
Control
Touch
n.s.
Average rosette radius(mm)
Days after touch
Double blinded experiment
n.s.
28.4±0.2
28.8±0.2
Days to bolting
Control
Touch
n = 68
n = 69
Control
Touch
Bolting plants(%)
Days after sowing
Control
Touch
n.s.
Average rosette radius(mm)
Days after touch

**Figure S11. Touch responses of soil-grown *arc3pmi4* Arabidopsis to automated human hair touch.**

The comparison of bolting plants grown in soil and touched by automated human hair. The above-left panel is the average bolting percentage from all biological experiments. The box and whisker plots of bolting days were analyzed for each biological experiment. The numbers of the total individual plants (n) and means ± SE from 3 biological replicates and one double-blinded experiment are annotated under and above the plots, respectively. The two right-above panels are photographs of a representative of untouched (control) and touch-treated plants (touch) for bolting and plant size. The number of bolting plants was the average bolting number calculated by three people. Trend chart plots which are the average rosette radius over the touch period (touch day) are shown in the right panels. Statistical test was performed employing student's t-test. The significance of $p < 0.05$, $p < 0.01$, and $p < 0.001$ for pairwise t-test are shown as *, **, and ***, respectively. T-DNA insertion mutant *arc3pmi4* was applied in this figure. This figure is related to Fig. 4.

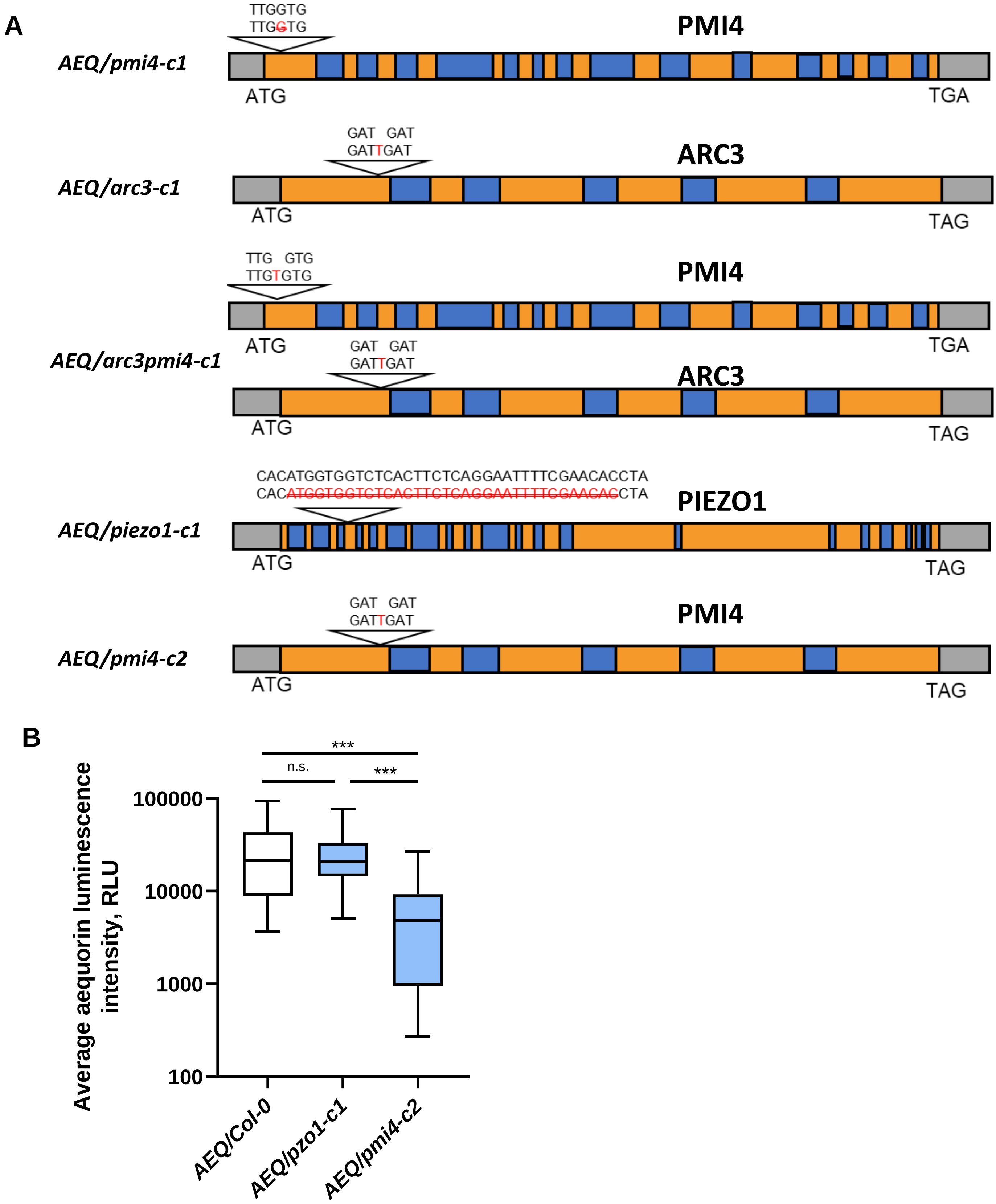


**Fig. S12. CRISPR generated mutant on *AEQ* background.** (A) Sequencing result of all CRISPR-Cas9 generated $Ca^{2+}$ reporter mutant used in this study. 'ATG' indicates the start codon while 'TAG'/'TGA' indicates the stop codon of each gene. Upper string sequences are the original genomic sequence, and the lower string sequences are the edited sequences. Blue segments represent intron and orange segments represent exon. (B) Comparison of peak intensity value among the tested plants. n = 25, 17, 22 for three genotypes, respectively. This figure is related to Fig. 5.

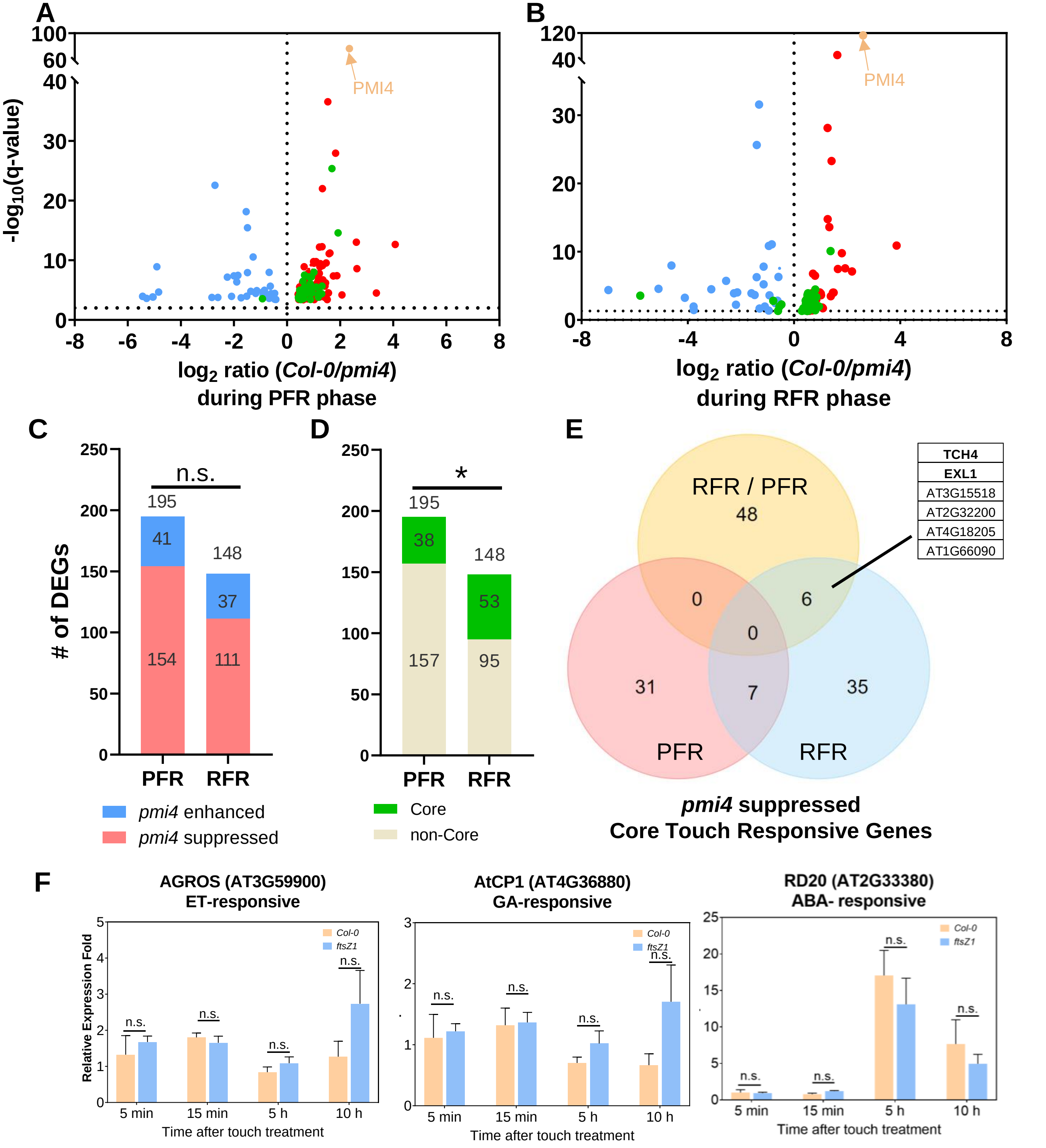


**Figure S13. Transcriptome analysis of *pmi4* mutant and wild-type *Col-0* plants under Prompt Force Response (PFR) condition vs Rapid Force Response (RFR) conditions.** (A-B) A volcano plot showing the 195 DEGs in PFR phase (A) and 148 DEGS in RFR phase (B), related data can be found in Supplemental Table S3a, S3b. Green dots indicate the "core touch responsive genes". (C) Summarization of *pmi4*-suppressed ($log_2$ (*Col-0*/*pmi4*) > 0, q-value <0.05) or *pmi4*-enhanced ($log_2$ (*Col-0*/*pmi4*) < 0, q-value < 0.05) DEGs composition in PFR and RFR phases, fisher`s exact test was applied to compare the composition, n.s. indicates not significant. (D) Summarization of "core touch responsive gene" composition in PFR and RFR phase, fisher`s exact test was applied to compare the composition, * indicates p value <0.05. (E) A Venn diagram showing the comparison of the core touch responsive gene from either PFR, RFR, and RFR/PFR *pmi4* repressed DEG list. (F) Changes in the relative expression levels of *CML38*, *AGROS*, and *AtCP1* in *Col-0* and *pmi4* at different time points after touch treatment. Relating data can be found in Supplemental Table S3d. The mRNA levels were quantified using RT-qPCR by the $2^{-\Delta\Delta CT}$ method. Mean ±SD of four biological repeats are shown. Student's t test was used to analyze the statistics: * indicates p value <0.05. T-DNA insertion mutant *pmi4* was applied in this figure. This figure is related to Fig. 6.

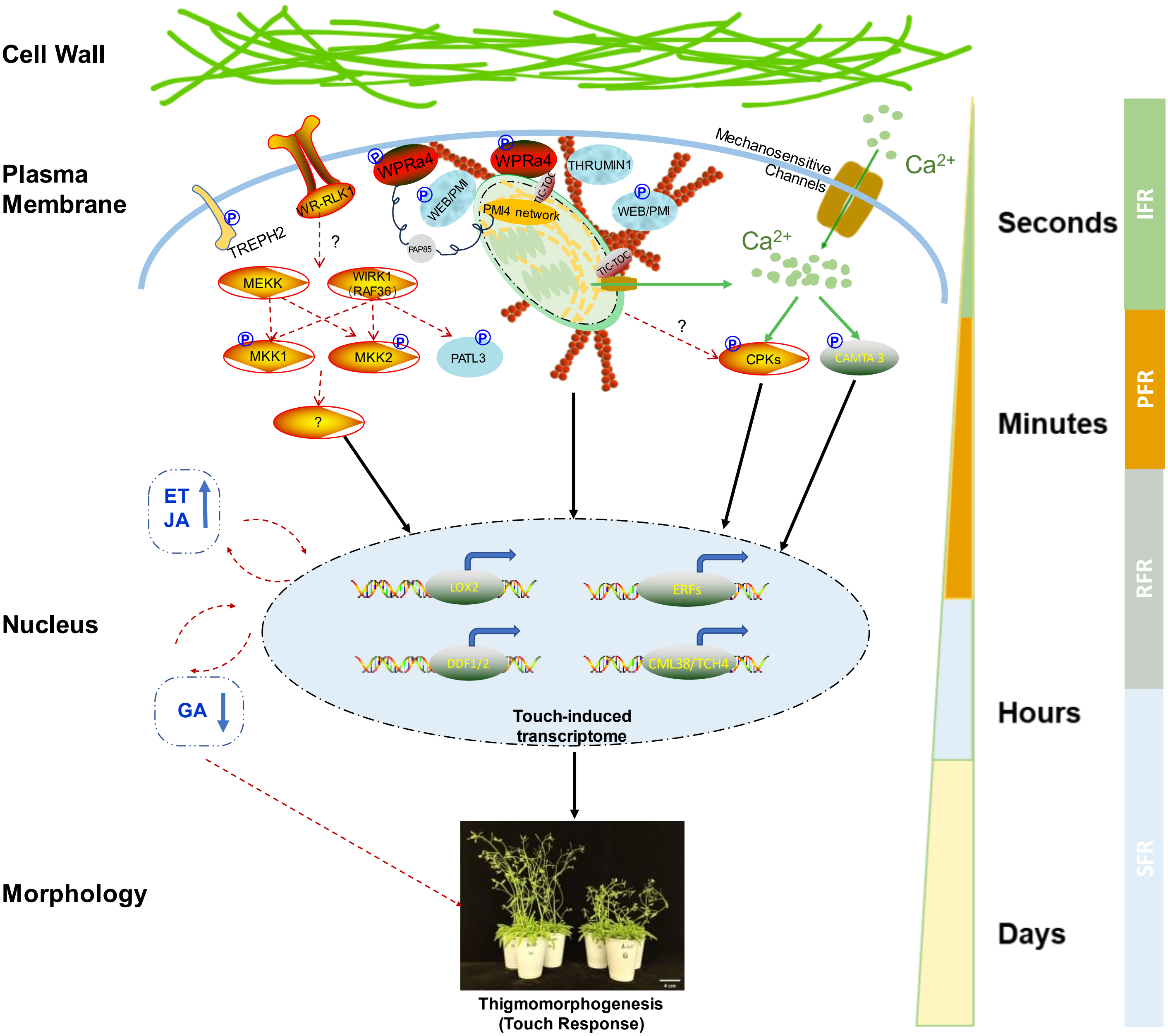


**Figure S14. A working module of WPRa1 and chloroplast-related touch-responsive gene expression.** Mechano-stimulation starts from physical pressure on cell walls, which then triggered the remodeling of the plasma membrane and cytoskeleton system. Temporal development with time scale and different force-responsive phases were classified (see Material and Methods). A possible linkage between WPRa4 and PMI4 network through TIC-TOC complex (PL data from this study) or PAP85 (crosslinking data from (Dai et al., 2023)). Plastid senses the mechano-stimulation through the intrinsic cytoskeleton protein PMI4 forming Z-ring and interacts with the cellular cytoskeleton system for relocation response. On the other hand, membrane $Ca^{2+}$ channels and plastid $Ca^{2+}$ waves were activated, promoting rapid increase of $Ca^{2+}$ concentration, including calcium-dependent protein CPKs activation (not shown) and calmodulin activation. Phosphorylation events from both possible CPKs and known MEKK-MKKs-WIRK1 signaling pathways were then activated, which regulates the downstream activation transcription factors into the nucleus, and promotes touch-responsive gene expression,e.g. TCHs, ERFs, et al. Phytohormone JA and ET were upregulated while the GA was down-regulated several hours later after the touch treatment. Long-term supply of mechano-stimulation causes evident alteration in thigmomorphogenesis.
